# LongAllele: a joint inference framework for allele-specific analysis on long-read bulk and single-cell RNA sequencing

**DOI:** 10.64898/2026.05.05.722992

**Authors:** Zhuoran Xu, Kai Wang

## Abstract

Allele-specific analysis from RNA-seq is a powerful approach to characterize *cis*-regulatory effects. However, existing methods remain limited in both haplotype inference and allelic testing. Their haplotype-inference workflows separate variant calling, haplotype phasing, and read-haplotype assignment into sequential steps, failing to fully exploit within-read single-nucleotide variant (SNV) linkage information and propagating errors into downstream allelic analysis. At the testing stage, they ignore non-phasable reads lacking heterozygous SNVs, biasing calls and inflating false positives, and remain incomplete across gene-, isoform-, and local-event-level variant effects. Here, we present LongAllele, a statistical framework that employs an expectation-maximization algorithm to jointly infer heterozygous variants, haplotype structure, and read-haplotype assignments from long-read bulk and single-cell RNA sequencing. LongAllele further introduces phasability-aware testing that explicitly accounts for non-phasable reads, avoiding inflated false-positive calls when haplotype information is incomplete. It also enables comprehensive allelic testing across gene-level allele-specific expression (ASE), isoform-level allele-specific transcript usage (ASTU), and local-event-level haplotype-associated exon and junction usage (HAEU and HAJU), providing a multi-scale view of *cis*-regulation across biological contexts. We applied LongAllele to long-read RNA-seq datasets spanning GTEx (multi-tissue bulk), peripheral blood mononuclear cells (single-cell), human hippocampus (single-nucleus), and human cortex from two Alzheimer’s disease (AD) case-control cohorts (bulk, Oxford Nanopore and PacBio). LongAllele consistently revealed greater context dependence in expression-level than isoform-level allelic regulation across tissues, cell types, and disease states, and pinpointed high-impact regulatory variants including rare splice-site mutations missed by standalone variant callers. It further showed that purifying selection constrains allelic imbalance at both gene and isoform levels and resolved AD-associated variant effects in individual transcriptomes across long-read platforms.

## 1 Introduction

Interpreting the functional impact of genetic variation is a central challenge in human genetics. By leveraging genetic and phenotypic differences across individuals in a population, genome-wide association studies (GWAS) and quantitative trait locus mapping, including expression QTLs (eQTLs), splicing QTLs (sQTLs) and methylation QTLs (meQTLs), have become powerful tools for linking genetic variants to disease risk [1] and quantifiable molecular phenotypes [2–6]. Notably, the majority of disease-associated variants identified by these studies reside in non-coding regions and are thought to act through *cis*-regulatory mechanisms [7, 8], yet interpreting their functional consequences remains difficult. While these population-level approaches have successfully identified genetic risk loci for many common diseases [9– 11], their reliance on between-individual variation necessitates large cohorts to adequately control for genetic and environmental confounders and to achieve sufficient statistical power. In contrast, allele-specific analysis compares molecular phenotypes between parental alleles within the same individual, inherently controlling for *trans*-acting and environmental confounders and enabling sensitive assessment of *cis*-regulatory variant effects [12, 13]. This within-individual design is particularly useful for rare variants, for which population-scale approaches are often underpowered [14–17]. Importantly, allele-specific analysis can resolve variant effects across multiple molecular scales. At the gene level, allele-specific expression (ASE) captures total-expression imbalance between parental haplotypes, whereas allele-specific transcript usage (ASTU) captures haplotype-dependent isoform shifts that may be invisible from gene-level analysis [2, 18, 19]. Local event-level analyses, including haplotype-associated exon usage (HAEU) and haplotype-associated junction usage (HAJU), further aid variant interpretation by resolving isoform-level imbalance into transcript-structural changes.

Realizing this multi-scale view requires resolving haplotype structure from heterozygous single-nucleotide variant (SNV) alleles and assigning reads to parental haplotypes. Although short reads can test local SNV associations with exon or splice-junction usage, linking SNVs within genes for haplotype-resolved analysis remains a fundamental limitation. Exonic heterozygous SNVs occur at only approximately 0.61 per kilobase across autosomal coding regions in GIAB HG001 [20], leaving few short reads that span multiple sites for phasing, with alternative splicing further compounding this challenge in RNA-seq [13, 21]. External haplotype information from population-level statistical phasing can partially address this challenge, but is largely restricted to common variants [22–24]. Individual-specific phasing from paired whole-genome sequencing (WGS) avoids this restriction [25, 26] but requires multi-omics profiling that is not always available. Even when SNV haplotypes are known, short-read allelic inference remains largely SNV-based, estimating gene-level haplotype imbalance from allelic proportions at individual variant sites. This SNV-based strategy can misrepresent gene-level haplotype expression when different isoforms cover distinct subsets of SNVs [13], and short reads cannot resolve full-length isoforms needed for transcript-level allelic analysis. By contrast, long-read RNA sequencing platforms, including Oxford Nanopore Technologies (ONT) and Pacific Biosciences (PacBio), address both limitations by capturing within-read SNV linkage for direct haplotype phasing while preserving full-length isoform identity for transcript-level analysis [21, 27]. These features support haplotype-resolved analysis of gene expression, transcript usage and local transcript events from RNA reads alone. Emerging long-read single-cell RNA sequencing technologies further extend haplotype-resolved isoform analysis to cellular resolution [28, 29], enabling analysis of how *cis*-regulatory effects on gene expression and transcript usage vary across cell types.

Despite these technological advances, computational methods for long-read allele-specific analysis remain underdeveloped (Table 1), often inheriting short-read assumptions [30] that limit both haplotype inference and allelic testing. In short-read workflows, haplotype inference is typically performed through sequential variant calling, haplotype phasing and read-haplotype assignment, whereas allelic testing often relies on SNV-level allele counts. Glinos et al. developed LORALS [25], one of the first methods for isoform-resolved allele-specific analysis using long-read RNA sequencing. LORALS requires paired WGS genotypes for DNA-guided haplotype inference and uses an SNV-based strategy for allelic testing. In this framework, long reads are used primarily to reduce haplotype-switching errors from HAPCUT2-based phasing [31], rather than to infer haplotype structure directly from SNV co-occurrence patterns within RNA reads. To eliminate the dependency on paired genotype data during haplotype inference, isoLASER [32] makes more direct use of long-read information by clustering reads into haplotype groups from SNV co-occurrence patterns. However, this weighted k-means clustering is heuristic and can be unstable when sparse SNV coverage and sequencing errors obscure allelic similarity among reads. For allelic testing, although isoLASER can test local exon-inclusion associations to distinguish *cis*-from *trans*-directed effects, it lacks gene- and isoform-level context for interpreting local events within broader allelic regulation. More recently, Huang et al. developed longcallR [33], which further improves genotype-free haplotype inference by replacing deterministic read clustering with a probabilistic read-based model, providing more stable read-haplotype assignments than heuristic clustering. Although longcallR enables gene-level ASE and local splice-junction testing, it does not provide isoform-level resolution for allelic analysis.

**Table 1.** Comparison of LongAllele with existing long-read RNA-seq allele-specific analysis methods.

|  |  | **LongAllele** | longcallR | isoLASER | LORALS |
| --- | --- | --- | --- | --- | --- |
| **Haplotype inference** | Paired genome data required | Optional | Optional | No | Yes |
|  | Variant calling | Joint EM | Standalone RNA-based caller | Standalone RNA-based caller | Standalone DNA-based caller |
|  | Haplotype phasing | Joint EM | Probabilistic read clustering | Heuristic k-means clustering | HAPCUT2 |
| **Allelic testing** | Testing strategy | Read-based | Read-based | Read-based | SNV-based |
|  | Phasability handling | Phasability-aware | Discard non-phasable | Discard non-phasable | Discard non-phasable |
|  | Resolution scales | ASE, ASTU, HAEU, HAJU | ASE, HAJU | HAEU | ASE, ASTU |
| **Data modality** | bulk / single-cell | bulk and single-cell | bulk only | bulk only | bulk only |

Still, all three methods inherit short-read-based sequential pipelines for haplotype inference and do not fully utilize within-read SNV co-occurrence information from long-read RNA-seq. They separate variant calling, haplotype phasing and read-haplotype assignment, so upstream errors can cascade into downstream allelic inference and testing. In long-read RNA-seq, however, heterozygous SNV evidence and haplotype structure could potentially inform each other, as SNV co-occurrence patterns can provide additional evidence for ambiguous variants while more accurate variant calls improve the resolution of haplotype structure. Beyond haplotype inference, existing long-read methods also remain limited in allelic testing. First, all existing methods assess haplotype imbalance exclusively from phasable reads that overlap heterozygous SNVs, implicitly assuming that these reads represent the full haplotype-level evidence for the tested gene. Since non-phasable reads provide no haplotype information and are therefore invisible to current models, ignoring their contribution goes beyond the power loss recognized previously [34] and can bias allelic estimates or produce false-positive calls. Second, although long-read methods have enabled isoform-resolved allelic analyses, no existing method comprehensively tests variant effects across gene-, isoform- and local-event-level regulation. Moreover, all three methods were developed for bulk long-read RNA-seq and do not support cell-type-resolved allelic analysis, leaving unaddressed the *cis*-regulatory heterogeneity across cell types that is unlocked by emerging long-read single-cell technologies.

Here, we present LongAllele, a statistical framework for joint haplotype inference and phasability-aware, multi-scale allelic testing from long-read bulk and single-cell RNA sequencing. LongAllele uses an expectation-maximization (EM) algorithm to jointly infer heterozygous variants, haplotype structure and read-haplotype assignments, allowing SNV evidence and haplotype structure to refine each other directly from RNA reads. We further introduce phasability-aware allelic testing that explicitly accounts for non-phasable reads and reduces the risk of biased or overconfident calls when phasable reads are not representative of the tested gene. Building on SCOTCH [35] isoform annotations, LongAllele provides comprehensive allelic testing across gene-level ASE, isoform-level ASTU, and local haplotype-associated exon and junction usage (HAEU and HAJU), at both bulk and cell-type resolution. We performed comprehensive simulation studies to benchmark LongAllele against existing methods for variant calling, haplotype phasing, read-haplotype assignment and allelic testing across varying allelic proportions and phasability conditions. We also applied LongAllele to long-read RNA-seq datasets spanning GTEx multi-tissue bulk samples, peripheral blood mononuclear cell (PBMC) single-cell data, human hippocampus single-nucleus data, and an Alzheimer’s disease case-control cohort, characterizing haplotype-resolved *cis*-regulatory effects across molecular, cellular and disease contexts. Overall, these analyses establish LongAllele as a unified framework for interpreting haplotype-resolved variant effects from bulk and single-cell long-read RNA sequencing.

## 2 Results

### 2.1 Overview of the LongAllele framework

LongAllele takes as input a genome-aligned BAM file from long-read RNA sequencing (bulk or single-cell), optionally with paired WGS genotypes, and produces haplotype-aware count matrices at both gene and isoform levels (Fig. 1). The framework combines read-to-gene and read-to-isoform mappings generated by SCOTCH [35] with candidate heterozygous SNVs identified through direct pileup (Fig. 1b). When paired genotype data are available, candidate SNVs are restricted to known heterozygous sites. Otherwise, candidates are filtered by heterozygous probability, homopolymer context, variant clustering, known A-to-I RNA editing sites [36], and an optional XGBoost classifier [37]. The filtered variants and read alignments are encoded into a read-variant profile matrix *r*, where *r*_*ij*_ ∈ { ref, alt, other } records the allele each read carries at each SNV, paired with a position-specific error rate matrix *π* derived from base quality (Phred) scores. The core of LongAllele is an EM algorithm that jointly infers latent variables of read-haplotype assignments *I*_*i*_ ∼ Bernoulli(*α*_*A*_) and SNV heterozygosity status 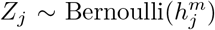, while iteratively re-estimating the haplotype-A proportion *α*_*A*_, SNV heterozygosity probability *h*^*m*^, and haplotype-A mapping probability *h* (Fig. 1b). This joint formulation leverages SNV co-occurrence patterns across reads to distinguish true heterozygous variants from sequencing noise, while accurate variant calls reciprocally improve haplotype phasing.

**Fig. 1.**
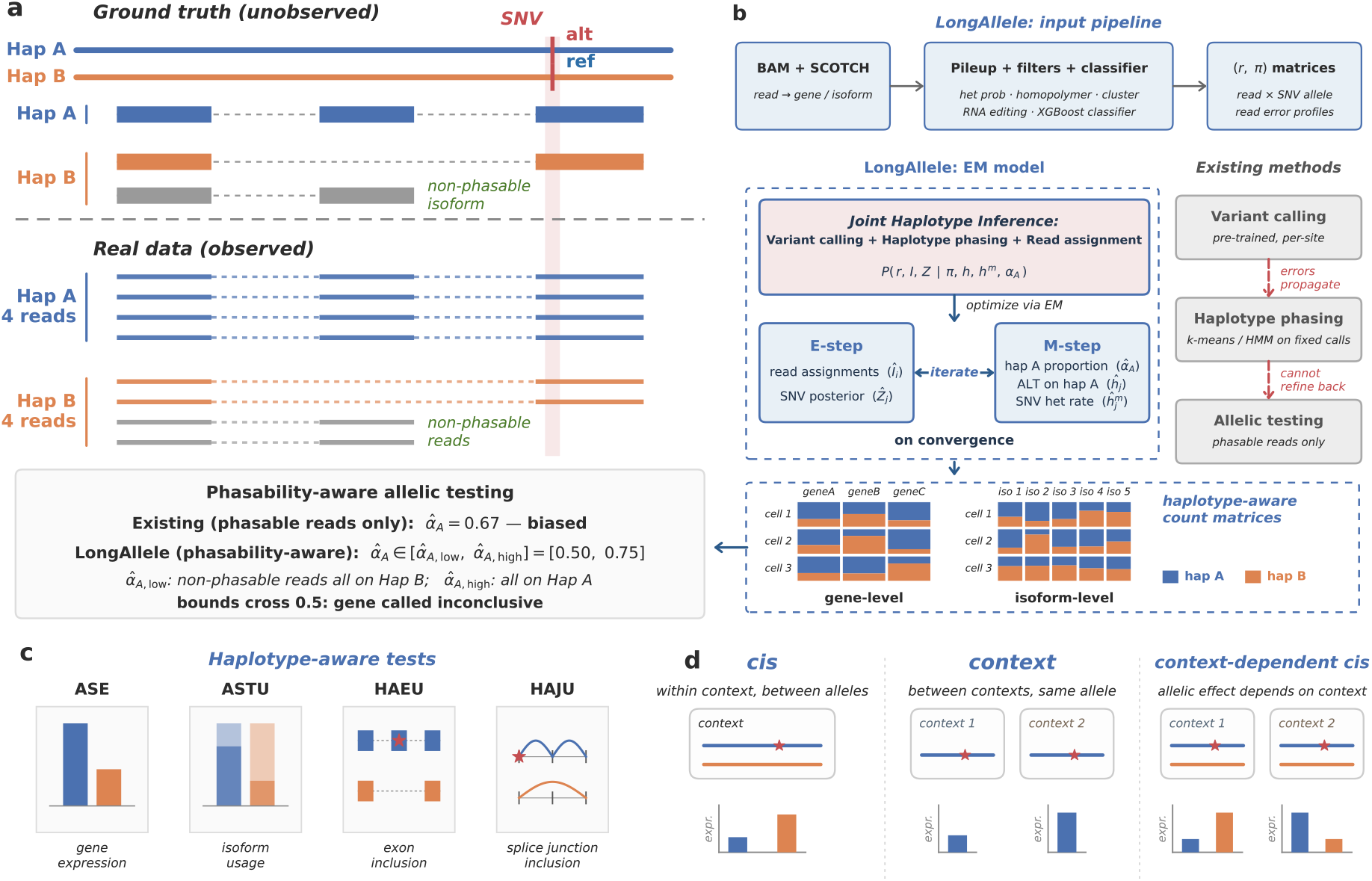
Overview of the LongAllele framework for haplotype-resolved allele-specific analysis from long-read single-cell RNA sequencing (lr-scRNA-seq). (**a**) Phasability-aware haplotype proportion inference on a single-gene toy example. *Top*, unobserved ground truth. A diploid genome carries one heterozygous SNV (red). One isoform (gray) does not cover the SNV and is non-phasable. *Bottom*, observed reads include phasable reads (blue/orange, covering the SNV) and non-phasable reads (gray). A phasable-only estimate 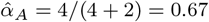 falsely declares allelic imbalance. LongAllele instead returns a phasability-aware interval 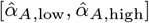 by assigning all non-phasable reads to Hap B or Hap A respectively, here [0.50, 0.75]. Because the interval spans 0.5, the gene is called inconclusive. (**b**) Joint inference framework of LongAllele. *Top*, inputs are a BAM file and SCOTCH read-to-gene and read-to-isoform mappings. Candidate SNVs are called by pileup and filtered by heterozygous probability, homopolymer context, variant clustering, RNA editing, and an optional XGBoost classifier, producing a read-by-SNV allele matrix and a per-position error matrix. *Left*, LongAllele jointly infers variant calls, haplotype phasing, and read-haplotype assignments through a single EM procedure. *Right*, sequential pipelines split these steps into independent stages where errors accumulate, and allelic tests consider only phasable reads. *Bottom*, LongAllele outputs haplotype-aware count matrices at both gene and isoform level (blue for Hap A, orange for Hap B). (**c**) LongAllele tests for allele-specific expression (ASE) and allele-specific transcript usage (ASTU) to quantify haplotype-level imbalance at gene and isoform levels, and identifies haplotype-associated exon usage (HAEU) and haplotype-associated junction usage (HAJU) to localize allelic effects to individual exons and splice junctions. (**d**) LongAllele distinguishes allelic effects from context effects. *Cis*-regulatory effects are measured as expression or transcript-usage differences between haplotypes within the same cellular context. Context effects are measured as differences across tissues, cell types or disease states for the same haplotype. Context-dependent *cis* effects arise when the magnitude or direction of allelic imbalance changes across contexts.

After convergence, LongAllele assigns each phasable read to a haplotype and generates haplotype-aware count matrices at gene and isoform levels, with fractional counts reflecting assignment uncertainty (Fig. 1b). We quantify haplotype-level imbalance in total gene expression as ASE, and in isoform composition as ASTU, extending these established concepts to the long-read haplotype-resolved setting. LongAllele tests for ASE using a likelihood ratio test and for ASTU using a chi-squared test on isoform-by-haplotype contingency tables, at both bulk and cell-type levels (Fig. 1c, Methods). Critically, reads not covering any heterozygous SNV (non-phasable reads) are explicitly modeled rather than discarded, with bounds on allelic estimates computed under most-balanced and most-imbalanced assignments of non-phasable reads (Fig. 1a). This yields a three-way classification of each gene as significant, non-significant, or inconclusive, avoiding overconfident calls under phasability uncertainty.

While ASE and ASTU quantify allelic imbalance at the gene and transcript levels, understanding the molecular mechanisms of *cis*-regulatory variants requires localizing their effects to individual exons and junctions. To this end, LongAllele identifies haplotype-associated exon usage (HAEU) and haplotype-associated junction usage (HAJU) (Fig. 1c), and tests for association between each event and nearby phased heterozygous SNVs. These associations can be further linked to non-coding regulatory variants through linkage disequilibrium (LD) when paired genotype data are available. Allelic regulation reflects the combined influence of *cis*-acting variants, cellular regulatory context, and interactions between them. Because two parental haplotypes are measured within the same cellular context, differences between haplotypes provide within-sample estimates of *cis*-regulatory effects (Fig. 1d). Conversely, differences across cellular contexts for the same haplotype capture context effects while holding genotype fixed. In addition, the magnitude or direction of the between-haplotype difference can vary across tissues, cell types, or disease states, revealing context-dependent *cis* regulation. By measuring haplotype-resolved allelic imbalance across contexts, LongAllele can help disentangle within-context *cis* effects, context effects, and context-dependent *cis* regulation.

### 2.2 LongAllele outperforms existing methods in variant calling, haplotype phasing, and allelic imbalance detection

To evaluate LongAllele’s performance in variant calling, haplotype phasing, and allelic imbalance detection, we simulated long-read RNA-seq data with realistic variant densities and expression levels estimated from the GIAB HG001 reference sample [20] and a long-read scRNA-seq PBMC dataset [35] (Methods). We simulated two phasability conditions on genes from chromosome 6, a fully phasable setting where every isoform covers at least one heterozygous SNV (464 genes), and a partially phasable setting where at least one isoform does not cover any heterozygous SNV (376 genes). For each phasability condition, we simulated three levels of allelic imbalance of gene expression with *α*_min_ ∈ { 0.1, 0.3, 0.5 }, representing strong, moderate, and no ASE. Of these, 160 multi-isoform genes were assigned ground-truth ASTU, with the rest being negative controls. We compared LongAllele against isoLASER [32] and longcallR [33], the two existing long-read RNA-seq methods that perform deep-learning-based variant calling followed by heuristic or probabilistic haplotype phasing. To benchmark performance when paired genotype data is available, we additionally included HAPCUT2 [31], a DNA-based haplotype phasing method that requires pre-called genotypes and is used by the LORALS pipeline [25], as well as LongAllele-Genotyped. Both were provided ground-truth SNV sites as input (Fig. 2).

**Fig. 2.**
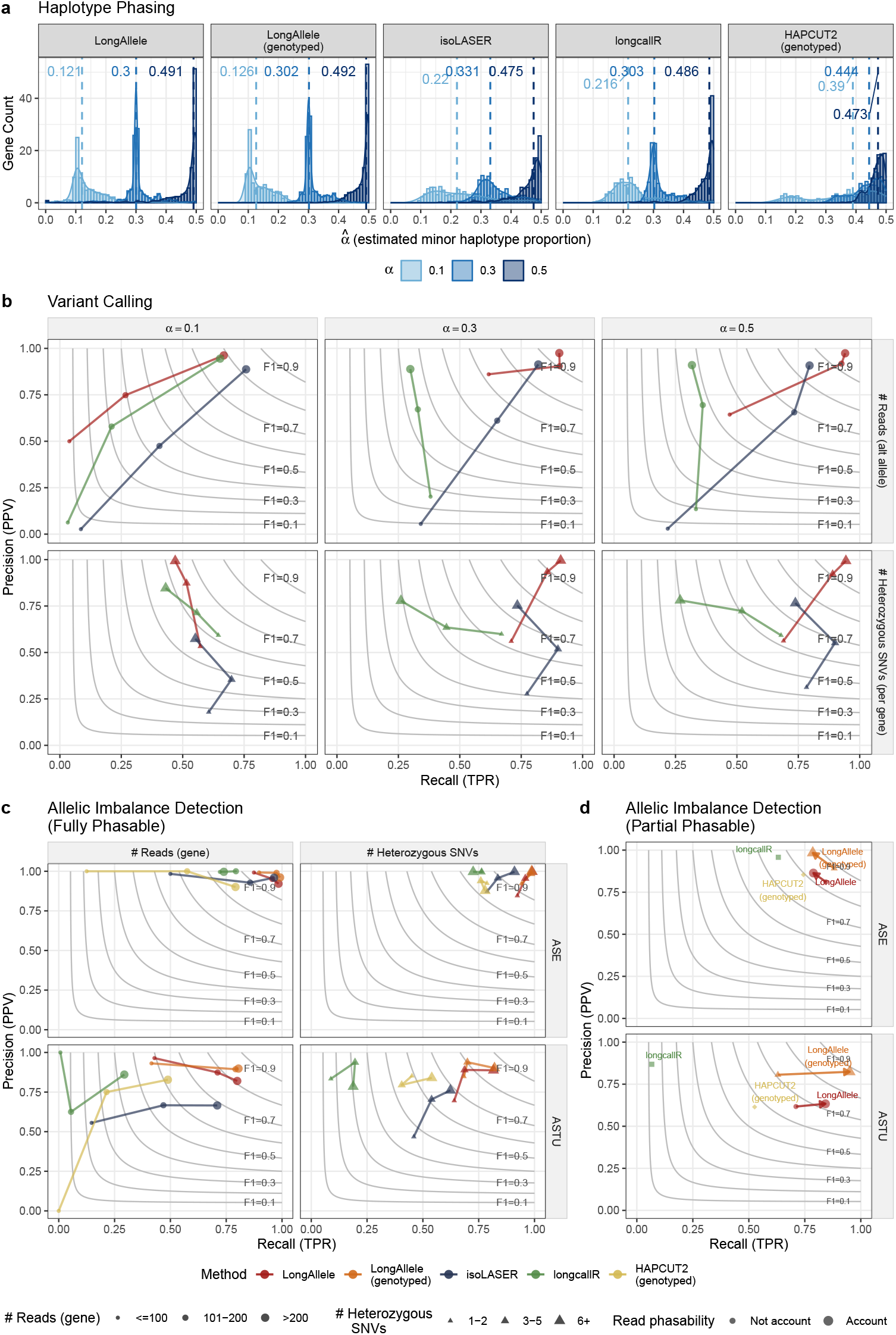
LongAllele outperforms existing long-read methods across haplotype phasing, variant calling, and allele-specific imbalance detection, with phasability accounting reducing false discoveries. (**a**)–(**c**) use the fully phasable setting (464 genes), and (**d**) uses the partially phasable setting (376 genes). Minor haplotype proportions (*α*_min_) were set to *α*_min_ = 0.1, 0.3, and 0.5 for both settings. LongAllele-Genotyped and HAPCUT2-Genotyped receive ground-truth variant sites as input. (**a**) Distributions of estimated 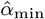 for all phased genes across methods. Dashed lines indicate median estimates. Numbers in parentheses indicate genes reported by each method. (**b**) Variant calling precision and recall, stratified by the number of reads supporting the alternative allele (top row) and the number of heterozygous SNVs per gene (bottom row). In (**c**) and (**d**), genes across *α*_min_ levels are pooled as the evaluation set (0.1 and 0.3 as positives, 0.5 as negatives). (**c**) Precision and recall for ASE (top) and ASTU (bottom) detection, stratified by the number of reads per gene and the number of SNVs per gene. (**d**) Precision and recall for ASE (top) and ASTU (bottom) detection in the partially phasable setting. Arrows point from performance without accounting for read phasability to performance with phasability accounted for. isoLASER was excluded because its phasing algorithm did not converge at *α*_min_ = 0.5.

We first compared estimated minor haplotype proportions 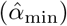 to their ground-truth values in the fully phasable setting (Fig. 2a). Since isoLASER and HAPCUT2 do not natively report per-gene allelic proportions and ASE significance, we derived these from their phased read assignments (Methods). Among all methods compared, LongAllele most accurately recovered true *α*_min_ across all settings, regardless of genotype availability, with median estimates closely matching the ground truth (LongAllele: 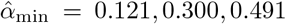, LongAllele-Genotyped: 0.126, 0.302, 0.492 for *α*_min_ = 0.1, 0.3, 0.5), indicating that accurate phasing can be achieved from RNA sequencing data alone. In contrast, isoLASER and longcallR overestimated *α*_min_ toward balanced proportions, with the bias most pronounced at *α*_min_ = 0.1 (median 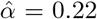 for both) and reduced at *α*_min_ = 0.3 (median 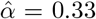 for isoLASER, 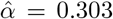 for longcallR), as neither method models the haplotype proportion *α* explicitly, leading to regularization toward balanced assignments when reads are weakly informative. This bias is even more pronounced for HAPCUT2, which produced estimates near 0.5 across all settings (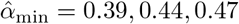 for *α*_min_ = 0.1, 0.3, 0.5) despite receiving true genotype information, reflecting the fundamental mismatch between DNA-based phasing methods that assume balanced allele representation and RNA-seq data where allelic proportions are determined by gene regulation. By contrast, LongAllele avoids this bias by learning *α* as an explicit parameter in the EM and feeding it back as a prior during read-haplotype assignment, so that the inferred proportion and assignments iteratively refine each other. Note that LongAllele phased over 99% of genes (460 of 464) across all *α*_min_ settings, whereas competing methods showed substantial gene loss, with isoLASER and HAPCUT2 phasing approximately 355 and 380 genes on average, and longcallR phasing only 332 genes (72%) at *α*_min_ = 0.1 due to reduced sensitivity to minor-haplotype SNVs at low allele fractions (Table S1).

We further evaluated variant calling performance across allelic imbalance settings for LongAllele, isoLASER, and longcallR (Fig. 2b). Performance improved for all methods as *α*_min_ increased toward balanced proportions. At *α*_min_ = 0.1, all methods performed similarly, reflecting the inherent difficulty of detecting heterozygous SNVs under highly skewed allele fractions. At *α*_min_ = 0.3 and 0.5, where both haplotypes contribute substantially to read coverage, LongAllele achieved higher F1 scores across read-depth strata, particularly at low alternative allele counts, by leveraging haplotype-level co-occurrence during variant calling. In contrast, longcallR exhibited high precision but lower recall under these settings, while LongAllele maintained both. Across genes with varying numbers of heterozygous SNVs, LongAllele’s variant calling performance increased with more SNVs per gene, as more variants strengthen the haplotype linkage signal used by joint inference. In contrast, isoLASER and longcallR did not exhibit this trend, as their pre-trained variant callers assess each candidate site independently without leveraging cross-variant linkage.

Having established differences in haplotype phasing and variant calling, we next assessed how these propagate to downstream ASE and ASTU detection by pooling genes across *α*_min_ levels (0.1 and 0.3 as positives, 0.5 as negatives), with each method evaluated on its own reported gene set (Fig. 2c, Fig. S1). For ASE detection, all methods achieved high precision, with performance primarily determined by recall, which increased with gene-level read coverage. At high coverage (*>*200 reads per gene), LongAllele and LongAllele-Genotyped achieved recall above 0.98, followed by isoLASER (recall = 0.96), whereas longcallR (recall = 0.74) and HAPCUT2 (recall = 0.79) showed substantially lower recall. This advantage persisted at lower coverage (≤ 100 reads), where LongAllele maintained recall of 0.87 while isoLASER dropped to 0.50. LongAllele also showed a clear dependence on the number of heterozygous SNVs per gene, with performance improving as SNV count increased and reaching an F1 score of 0.99 at six or more SNVs, reflecting the benefit of stronger haplotype linkage for joint inference of allelic imbalance. This pattern was absent in the other methods, with longcallR’s recall remaining around 0.72 to 0.76 regardless of heterozygous SNV count. ASTU detection proved more challenging than ASE, with larger performance gaps across methods. Because isoLASER, longcallR, and HAPCUT2 do not directly perform isoform-level allelic testing, we used their closest available outputs as proxies (Methods). At high coverage (*>*200 reads), LongAllele (F1 = 0.81) and LongAllele-Genotyped (F1 = 0.85) substantially outperformed isoLASER (F1 = 0.69), HAPCUT2 (F1 = 0.62), and longcallR (F1 = 0.44) (Fig. 2c). longcallR exhibited particularly low recall across all coverage levels, dropping to near zero at ≤ 100 reads (F1 = 0.01), consistent in part with the limited sensitivity of junction-level tests, which may fail to capture isoform shifts that do not alter individual splice junction usage. LongAllele’s ASTU detection also improved with increasing numbers of heterozygous SNVs per gene, with F1 increasing from 0.67 at one to two SNVs to 0.85 at six or more SNVs. This dependence on SNV count was weak or absent in other benchmark methods.

In practice, a substantial fraction of reads do not overlap any heterozygous SNV and cannot be assigned to a haplotype. Existing methods discard these non-phasable reads, risking overconfident calls when phasable coverage is sparse. We evaluated this scenario using the partially phasable setting, where LongAllele’s phasability-aware testing improves reliability by accounting for the unobserved reads (Fig. 2d, Fig. S2). For ASE, precision increased from 0.81 to 0.86 for LongAllele and from 0.89 to 0.98 for LongAllele-Genotyped, with overall F1 nearly unchanged (0.83 to 0.82 and 0.89 to 0.87, respectively), as 25% and 37% of genes were reclassified as inconclusive. For ASTU, phasability adjustment improved both precision and recall, with F1 increasing from 0.66 to 0.72 for LongAllele and from 0.71 to 0.88 for LongAllele-Genotyped. Notably, phasability adjustment yielded a larger gain for LongAllele-Genotyped than for LongAllele, whereas the two performed similarly in the fully phasable setting. This suggests that variant calling errors interact with incomplete read phasability to affect downstream allelic imbalance inference, and that providing known genotypes mitigates this source of noise. In contrast, longcallR and HAPCUT2, which do not account for phasability, showed substantially lower performance, while isoLASER failed to converge under balanced conditions (Tables S2-S5). By jointly modeling SNV zygosity and co-occurrence patterns, LongAllele achieves more accurate variant calling and haplotype phasing than sequential pipelines, translating directly into improved ASE and ASTU detection. Explicit accounting for non-phasable reads further improves reliability by flagging ambiguous cases as inconclusive rather than producing false calls.

### 2.3 LongAllele reveals context-dependent *cis* regulation in GTEx long-read bulk RNA-seq

Building on the robust performance observed in simulations, we further evaluated LongAllele on bulk long-read RNA-seq from seven GTEx donors, each profiled in two tissues across nine distinct tissue types (14 donor-tissue samples in total). Using exonic heterozygous SNVs from paired WGS as ground truth, LongAllele-Genotyped achieved F1 score of 0.856 for variant calling (Fig. 3a, Table S8), setting an intrinsic upper bound for genotype-free performance, as the EM strategically downweights SNVs with weak haplotype linkage signal regardless of known genotype status. Without genotype input, LongAllele and longcallR achieved near-identical F1 scores of 0.709 and 0.707, respectively, demonstrating comparable variant calling performance from RNA-seq data alone. Because no ground truth exists for ASE, we measured ASE concordance rate between each method and LORALS on genes where both produced conclusive calls. Across 14 samples, all three methods achieved over 90% concordance with LORALS (LongAllele-Genotyped 93.4%, LongAllele 92.1%, longcallR 90.1%, Table S7), confirming that genotype-free approaches produce ASE calls comparable to genotype-dependent methods. Quantitatively, haplotype fraction estimates 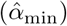 derived from LongAllele and longcallR showed significantly higher concordance (Pearson *r* = 0.62) than those between LongAllele-Genotyped and LORALS (*r* = 0.14, both *p <* 0.001, Fig. S3b). This divergence stems from the fact that both LongAllele and longcallR utilize haplotype linkage information from full-length reads, whereas LORALS relies on global proportions inferred from local HAPCUT2 phase blocks (Fig. S3c), inheriting DNA phasers’ bias toward balanced allelic proportions.

**Fig. 3.**
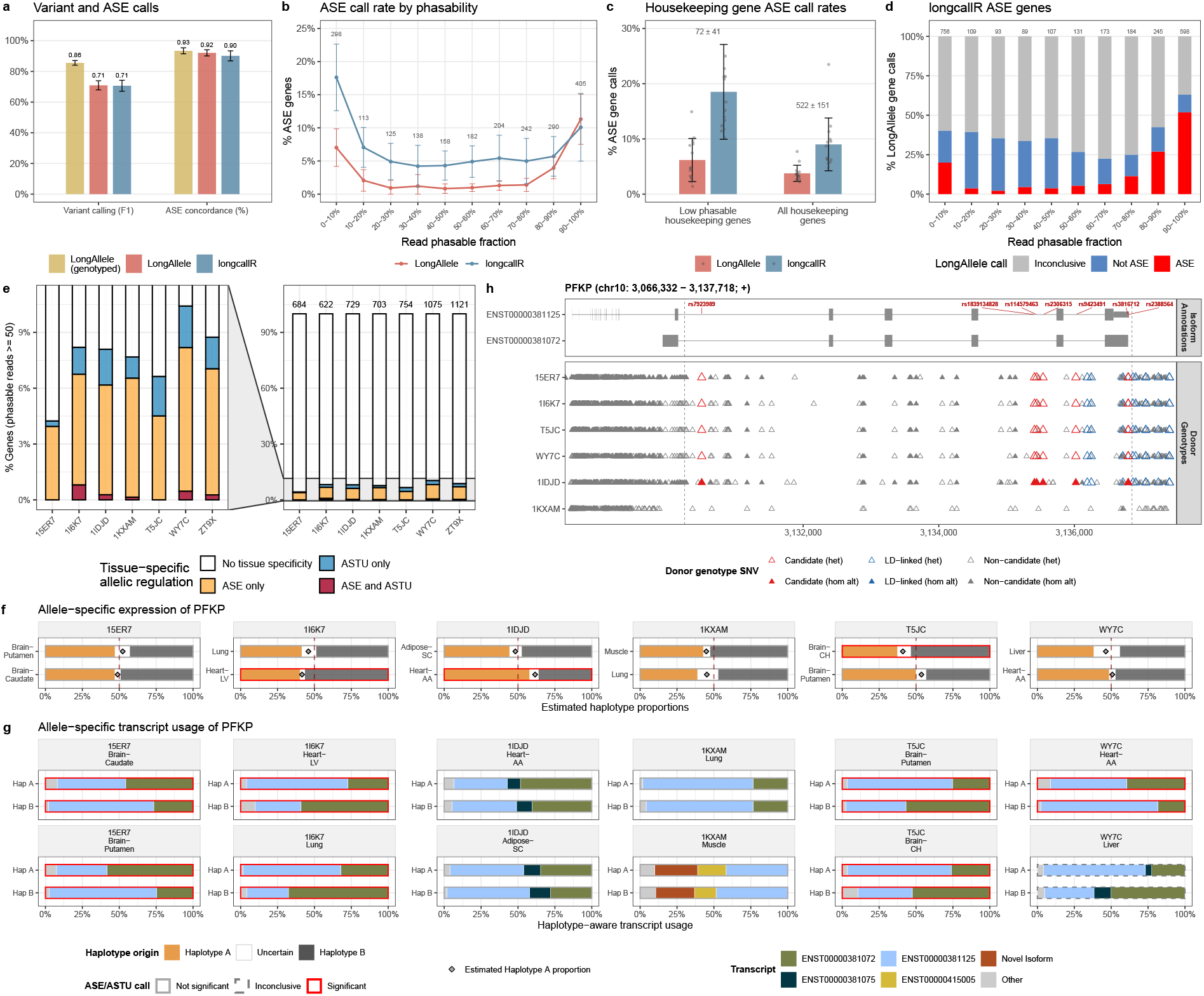
Allele-specific analysis of GTEx bulk long-read RNA-seq with multi-tissue donors. LongAllele provides more reliable ASE calls with phasability-aware testing and helps disentangle within-tissue *cis* effects, tissue effects, and context-dependent *cis* regulation across tissues. (**a**) Left, F1 scores for variant calling performance. Exonic heterozygous SNVs from WGS are used as ground truth. Right, ASE concordance rate with LORALS across 14 tissue samples. Error bars indicate standard deviation across tissue samples. (**b**) Percentage of genes called ASE-significant by LongAllele and longcallR across levels of gene phasability. The x-axis represents read phasable fraction, calculated as the number of reads overlapping at least one WGS-verified exonic heterozygous SNV over total reads per gene. Numbers represent genes phased by both methods in each bin. (**c**) Percentage of housekeeping genes (HRT Atlas) called ASE-significant. Left, low-phasability housekeeping genes (0–10% phasable reads). Right, all housekeeping genes phased by both methods. (**d**) LongAllele’s assessment of longcallR ASE-significant genes, stratified by read phasable fraction. Colors indicate LongAllele’s call on each gene (ASE-significant, not significant, or inconclusive). (**e**) Tissue-specific allelic regulation across 7 donors (2 tissues each). Right, percentage of testable genes (≥ 50 phasable reads in both tissues) showing tissue-specific ASE only, ASTU only, both, or neither. Numbers above bars indicate the number of testable genes per donor. Left, zoomed view of tissue-specific categories. (**f**) Allele-specific expression of *PFKP* across 6 donors (2 tissues each). Stacked bars show estimated haplotype-A (orange) and haplotype-B (grey) proportions with uncertainty intervals. Diamonds indicate point estimates of 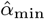. Borders represent ASE call significance. (**g**) Allele-specific transcript usage of *PFKP* across 6 donors (2 tissues each). Stacked bars show isoform proportions for each haplotype per donor-tissue. Borders represent ASTU call significance. (**h**) Candidate *cis*-regulatory SNVs for allele-specific isoform usage of *PFKP*. Top, transcript structures with candidate SNV positions (red labels), with 5 in shared introns and 2 at the 3’ end within a 9 bp differential sub-exon between the two isoforms. Bottom, donor genotype panel across the gene locus. Red triangles indicate 7 candidate SNVs associated with ASTU between ENST00000381125 and ENST00000381072, heterozygous in all 4 ASTU-positive donors (15ER7, 1I6K7, T5JC, WY7C) and homozygous in the remaining donors. Blue triangles indicate 10 deprioritized LD-linked variants (*r*^2^ *>* 0.9) with inconsistent genotype-phenotype patterns. In (**f**)–(**h**), haplotypes are aligned across donors based on shared SNVs.

To examine how non-phasable reads compromise ASE calling reliability on real data, we compared ASE call rates between LongAllele and longcallR across bins of phasable read fraction (Fig. 3b). We found that longcallR consistently called a higher proportion of genes ASE-significant than LongAllele at phasable fractions below 90%, with the gap narrowing at higher fractions and LongAllele slightly exceeding longcallR at 90-100%. This pattern suggests that LongAllele tends to draw ASE calls from genes with stronger haplotype evidence than longcallR. Notably, genes in the 0–10% phasability bin contain too few phasable reads to support confident allelic inference, yet LongAllele and longcallR called 7.0% and 17.6% of genes ASE-significant, respectively. In this regime, even a small number of false positive variant calls can overwhelm the limited set of true heterozygous sites, creating an apparent signal of phasability that leads to overconfident ASE calls. Further examination revealed that approximately a quarter of genes in this bin are housekeeping genes [38, 39], which are highly expressed but have low exonic heterozygous SNV density. Because housekeeping genes are predominantly biallelically expressed across cell types [40], they serve as an imperfect negative-control set, where the proportion called ASE-significant provides a relative proxy for each method’s false-positive rate (recognizing that some house-keeping genes may still be allele-imbalanced). Among these low-phasability housekeeping genes, longcallR called 18.5% as ASE-significant compared to 6.2% for LongAllele. This disparity persisted across all housekeeping genes phased by both methods (longcallR: 9.0% vs LongAllele: 3.8%, Fig. 3c), supporting the improved reliability of LongAllele through reduced false positive ASE calls. To further assess how phasability-aware testing affects ASE calls, we examined genes identified as ASE-significant by longcallR and evaluated LongAllele’s calls on the same set (Fig. 3d). Concordance increased monotonically with phasability, from 3% at 10-30% to 52% at 90-100%, showing that LongAllele retains longcallR’s ASE calls primarily when sufficient phasing evidence is available. Together, these results support that phasability-aware testing reduces likely false discoveries in real data, consistent with the simulation findings.

We next leveraged the multi-tissue design of GTEx data to examine within-tissue *cis* effects, tissue effects, and context-dependent *cis* regulation across tissues through haplotype-resolved inference. Tissue-specific regulation was defined as allelic imbalance that is significant in one tissue but not the other within the same donor. Among genes with ≥ 50 phasable reads in both tissues, most showed consistent allelic patterns across tissues on both gene and isoform levels, with only 4.2-10.4% of genes per donor displaying tissue-specific regulation (Fig. 3e). Among genes displaying tissue-specific regulation, ASE differences predominated over ASTU differences, indicating that ASE is more strongly modulated by tissue context while isoform usage remains relatively stable.

As a representative example, *PFKP* illustrates all three regulatory modes within a single gene (Fig. 3f,g). LongAllele phased *PFKP* in 12 of 14 donor-tissue samples across six donors, comparable to longcallR, whereas LORALS could test it in only one sample (T5JC Brain-CH, 36 reads), highlighting the greater read utilization of read-based than SNV-based testing. After aligning haplotypes across donors using shared heterozygous SNVs, three donors showed tissue-specific ASE, compared with one detected by longcallR. In these donors, significant allelic imbalance was present in one tissue but not the other, indicating that the *cis*-regulatory effect on total expression is modulated by tissue context. At the isoform level, several donors showed tissue-dependent changes in transcript usage without haplotype imbalance. For example, in 1KXAM, both haplotypes expressed ENST00000381125 and ENST00000381072 in lung but shifted to ENST00000415005 and a novel isoform in muscle. Because these changes affected both haplotypes equally, they were not detected as ASTU and instead reflected a tissue-context effect on transcript usage without detectable haplotype imbalance. By contrast, four of six donors showed significant ASTU, with haplotype A consistently switching from the canonical protein-coding transcript ENST00000381125 to ENST00000381072, a non-coding transcript with an alternative internal TSS and a 9 bp shorter 3’ UTR, indicating a *cis*-directed effect on isoform choice that is stable across tissue contexts. This pattern was observed in seven of eight donor-tissue samples, with the remaining sample inconclusive because of low read coverage.

To identify candidate *cis*-regulatory variants underlying ASTU in *PFKP*, we examined haplotype-associated exon usage (HAEU) and junction usage (HAJU) and linked these events to nearby heterozygous SNVs (Fig. 3h). Two SNVs (rs3816712 and rs2388564) within the structurally divergent region between ENST00000381125 and ENST00000381072 showed significant HAEU in all seven donor-tissue samples (phasability-adjusted *p <* 0.05). Using paired WGS genotypes, we found that these SNVs were in perfect linkage disequilibrium (*r*^2^ = 1.0) with five additional intronic variants, defining a seven-variant haplotype block. The alternative alleles of this block were consistently assigned to haplotype A, which favors ENST00000381072, and were heterozygous in all four ASTU-positive donors but homozygous in the two ASTU-negative donors. Structurally, rs3816712 and rs2388564 fall within the 9 bp differential sub-exon at the 3’ end and lie 1-2 bp from the proximal cleavage site of ENST00000381072 at chr10:3,136,793, which is supported by a canonical ATTAAA hexamer 10 nt upstream. Although these SNVs implicate allele-specific cleavage-site usage as the candidate mechanism for the 3’ UTR difference [41–43], they cannot account for the alternative internal TSS at the 5’ end. This 5’ structural difference may reflect allele-specific promoter usage driven by a co-inherited regulatory variant on the same LD block, or arise as a downstream consequence of the 3’ APA event through TSS-polyadenylation crosstalk. No other variants within ± 50 kb of the *PFKP* locus matched this genotype-phenotype pattern in the WGS data, making this haplotype block the leading candidate *cis*-regulatory element underlying the observed allele-specific isoform usage, with the 5’ difference likely reflecting indirect coupling to this APA event.

### 2.4 LongAllele resolves cell-type-specific allelic regulation masked in bulk from PBMC lr-scRNA-seq

Although bulk analysis can reveal allelic regulation at both gene and isoform levels, it inherently averages signals across heterogeneous cell populations, potentially masking cell-type-specific effects. To resolve allelic regulation at single-cell resolution, we applied LongAllele to Nanopore lr-scRNA-seq data from PBMCs of two healthy donors (Fig. 4). We identified ASE genes consistently detected across both donors within each cell type, restricting to genes with ≥ 20 phasable reads in at least two cell types to minimize confounding by low expression. While some ASE genes were significant in both bulk and individual cell types, a substantial subset showed significance exclusively at the cell-type level (red bars, Fig. 4a). These results indicate that bulk analysis can dilute cell-type-specific allelic signals and, in some cases, cancel them entirely when allelic directions are opposite.

**Fig. 4.**
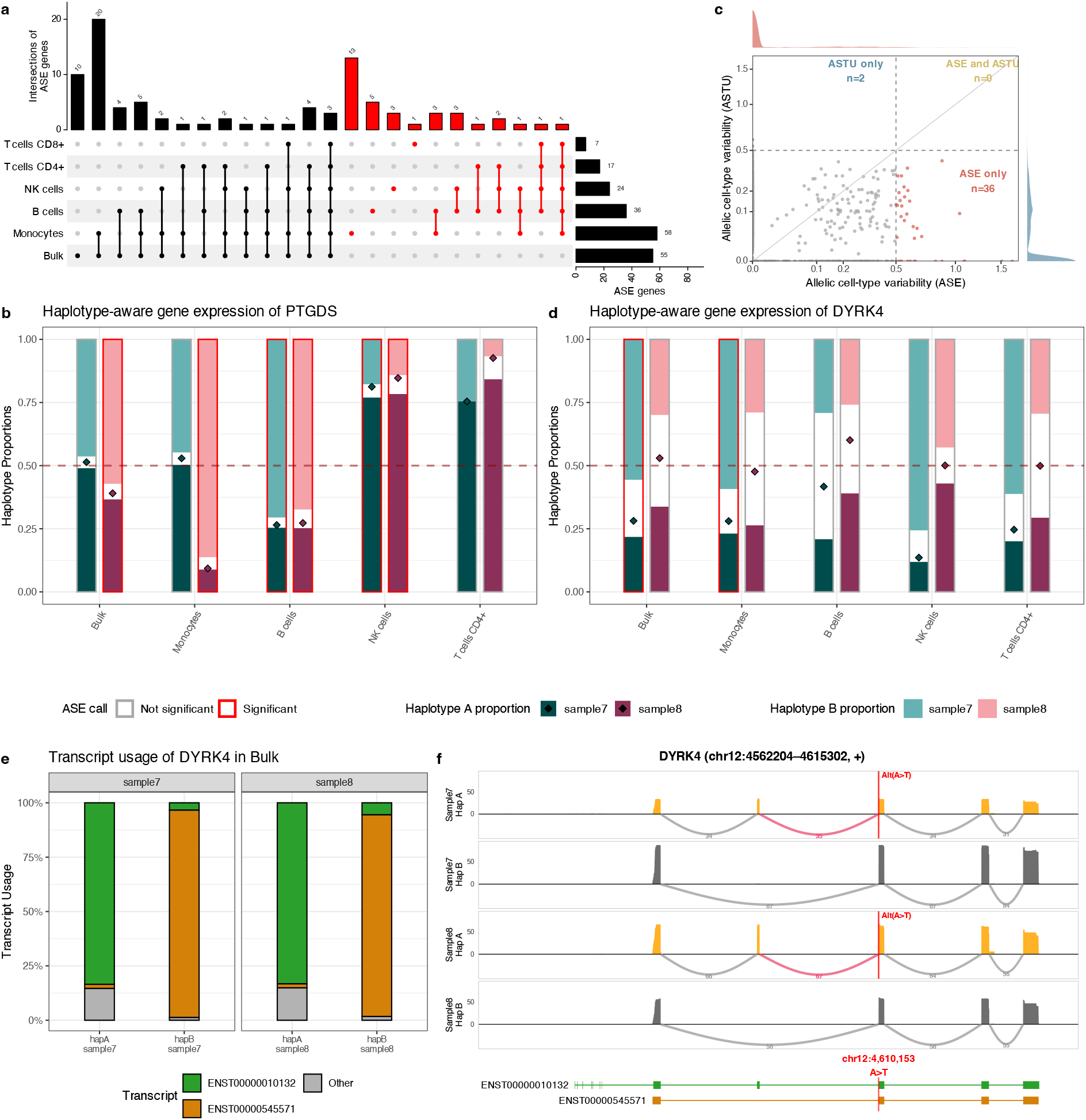
Cell-type-resolved allele-specific analysis in PBMC long-read single-cell RNA-seq. LongAllele uncovers cell-type-specific allelic regulation hidden in bulk and reveals greater cell-type variability for gene expression than for isoform usage in PBMC. (**a**) UpSet plot of ASE genes shared across both donors per cell type. Only genes with ≥ 20 phasable reads in at least two cell types are included. Red bars and points indicate genes called ASE-significant in at least one cell type but not in bulk. (**b**) Haplotype-aware gene expression of *PTGDS* across cell types. (**c**) Allelic cell-type variability (ACTV) scatter plot. Each dot represents one gene-donor pair. The x-axis shows ASE ACTV and the y-axis shows ASTU ACTV, measuring the variability of allelic effects across cell types for gene expression and isoform usage, respectively. Dashed lines indicate substantial cell-type dependence at ACTV = 0.5. (**d**) Haplotype-aware gene expression of *DYRK4* across cell types. In (**b**) and (**d**), stacked bars show estimated haplotype-A (dark), haplotype-B (light), and uncertain (white) proportions, diamonds indicate point estimates of haplotype-A proportion 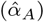 and red borders indicate significant ASE. (**e**) Haplotype-resolved transcript usage of *DYRK4* in bulk. (**f**) Haplotype-resolved sashimi plot of *DYRK4*. Red vertical line marks the position of a heterozygous variant (chr12:4,610,153, A*>*T) at the splice acceptor site. Arcs represent splice junctions with read counts, the pink arc highlights the SNV-proximal junction (chr12:4,607,327-4,610,154) directly affected by the variant. The differentially included exon spans chr12:4,607,327-4,607,387 (61 bp).

For example, *PTGDS* (prostaglandin D2 synthase), involved in immune modulation and expressed across multiple blood cell types, showed weak and inconsistent ASE at the bulk level (Fig. 4b, Fig. S4a). Sample 7 was essentially balanced 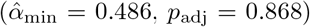, whereas sample 8 showed only moderate imbalance (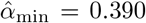, *p*_adj_ = 4.6 × 10^−4^). By contrast, cell-type-resolved analysis revealed not only clear ASE but also a reversal in allelic direction across cell types. In both donors, haplotype B was dominant in B cells, whereas haplotype A was dominant in NK cells (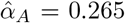 and 0.273 in B cells; 0.812 and 0.847 in NK cells). This opposing pattern explains why the bulk-level signal appears attenuated or inconsistent across donors. The reproducible direction switch across unrelated donors further suggests a shared *cis*-regulatory mechanism whose effect depends on cell-type context. LongAllele further identified rs6926 (chr9:136,981,689 A*>*C) in the 3’ UTR as the only shared heterozygous variant on haplotype A in both donors. This variant overlaps an ENCODE-annotated proximal enhancer [44] (cCRE EH38E3920662), is a known eQTL for *PTGDS* in blood [45], and has also been reported to drive allelic imbalance in heart tissue [46], supporting a functional *cis*-regulatory role across distinct tissue and cell-type contexts. Yet in prior bulk allelic-imbalance analyses, only 3% of heterozygous individuals showed significant imbalance for *PTGDS*, often with inconsistent effect directions [46]. A similar inconsistency is apparent across studies compiled in the eQTL Catalogue [45], supporting the view that bulk averaging obscures a structured, cell-type-specific regulatory pattern.

Beyond expression-level effects, we asked whether isoform-level allelic regulation also varies across cell types. To quantify this systematically, we defined allelic cell-type variability (ACTV), which measures the variability of *cis*-regulatory effects across cell types, with higher values indicating stronger cell-type dependence and lower values indicating more constitutive regulation (Methods). Genome-wide, *cis*-regulatory effects on gene expression showed substantially greater cell-type variability than those on isoform usage. Among genes with substantial cell-type dependence (ACTV ≥ 0.5), 36 showed high ASE variability compared with only 2 showing high ASTU variability, and no genes showed high variability in both (Fig. 4c). These results indicate that *cis*-regulatory effects on gene expression are more strongly shaped by cell-type context, whereas those on isoform usage remain comparatively constitutive. *PTGDS*, which showed strong cell-type-dependent ASE but no ASTU in any cell type, exemplifies this pattern (Fig. S5a).

As a complementary example to *PTGDS, DYRK4* showed constitutive ASTU across cell types in both donors, even though no significant ASE was detected in sample 8 (Fig. 4d, Fig. S4b, Fig. S5b). Across both donors, haplotype A predominantly expressed ENST00000010132, whereas haplotype B predominantly expressed ENST00000545571 (Fig. 4e). These two isoforms differ mainly by inclusion of a single exon and its associated splice junctions, both of which were strongly haplotype-dependent (HAEU *p*_adj_ = 1.1 × 10^−14^, 7.6 × 10^−13^; HAJU *p*_adj_ = 6.3 × 10^−15^, 2.0 × 10^−17^). LongAllele further pinpointed a heterozygous variant (chr12:4,610,153, A*>*T) at the splice acceptor site of this exon as the candidate *cis*-regulatory driver (Fig. 4f). The alternative allele promoted inclusion of this exon, thereby favoring ENST00000010132 over ENST00000545571, a pattern evident at the raw-read level in the long-read data (*p* = 1.6 × 10^−24^ and 3.8 × 10^−26^). Importantly, it was independently confirmed in paired short-read RNA-seq data (*p* = 1.1 × 10^−70^ and 2.9 × 10^−80^; Fig. S6), with reads carrying the alternative allele almost exclusively supporting the exon-included isoform. Independent computational evidence further supported this variant as a splice-disrupting allele. SpliceAI predicted the variant to cause high-confidence acceptor loss (delta score = 0.91) [47], and MaxEntScan predicted near-complete loss of splice-site strength (8.8 → 0.4) [48]. More generally, CADD ranked the variant among the top 0.08% most deleterious variants genome-wide (PHRED = 31) [49]. Despite this convergent evidence, the variant is absent from gnomAD v4.1 in both raw and filtered call sets [50], likely because conventional variant callers systematically miss such sites at the calling stage due to their non-canonical splice context and AT-rich flanking sequence, whereas LongAllele recovers such variants through direct pileup with haplotype linkage information. These two annotated isoforms are also distinguished by 5’ exons with little coverage (Fig. 4f). Thus, although the splice-site variant is strongly linked to the local exon-inclusion event, the full 5’ isoform structure remains ambiguous, as SCOTCH may assign a 3’-matched read to ENST00000010132 as a 5’-truncated read, leaving a novel isoform unexcluded. Together, this case shows how LongAllele can resolve constitutive allele-specific splicing and identify plausible regulatory variants that are readily missed by bulk or conventional variant-calling approaches.

### 2.5 LongAllele reveals purifying-selection signatures of allelic regulation in human hippocampus

To further examine allelic regulation in a more cell-type-diverse tissue, we applied LongAllele to Nanopore single-nucleus RNA-seq (lr-snRNA-seq) of postmortem human hippocampus from six healthy donors [51]. We performed hierarchical clustering of samples based on ASE and ASTU effect sizes to assess how cell-type context modulates genotype-driven allelic regulation (Fig. 5b). Samples from neurons, glia, and progenitors grouped primarily by donor rather than cell type, indicating that individual genotype is the dominant determinant of the allelic landscape, whereas cell-type identity contributes a secondary layer of modulation. In contrast, vascular and immune cells formed a distinct cluster characterized by uniformly low ASE and ASTU effect sizes, consistent across donors. The regulatory architectures of ASE and ASTU also differed, with ASE appearing as discrete donor-specific gene modules and ASTU as more diffuse and pervasive patterns (Fig. 5b). This qualitative difference was confirmed by ACTV quantification, which showed greater cell-type variability for ASE than for ASTU (Fig. S7), consistent with the same pattern observed in PBMCs (Fig. 4c).

**Fig. 5.**
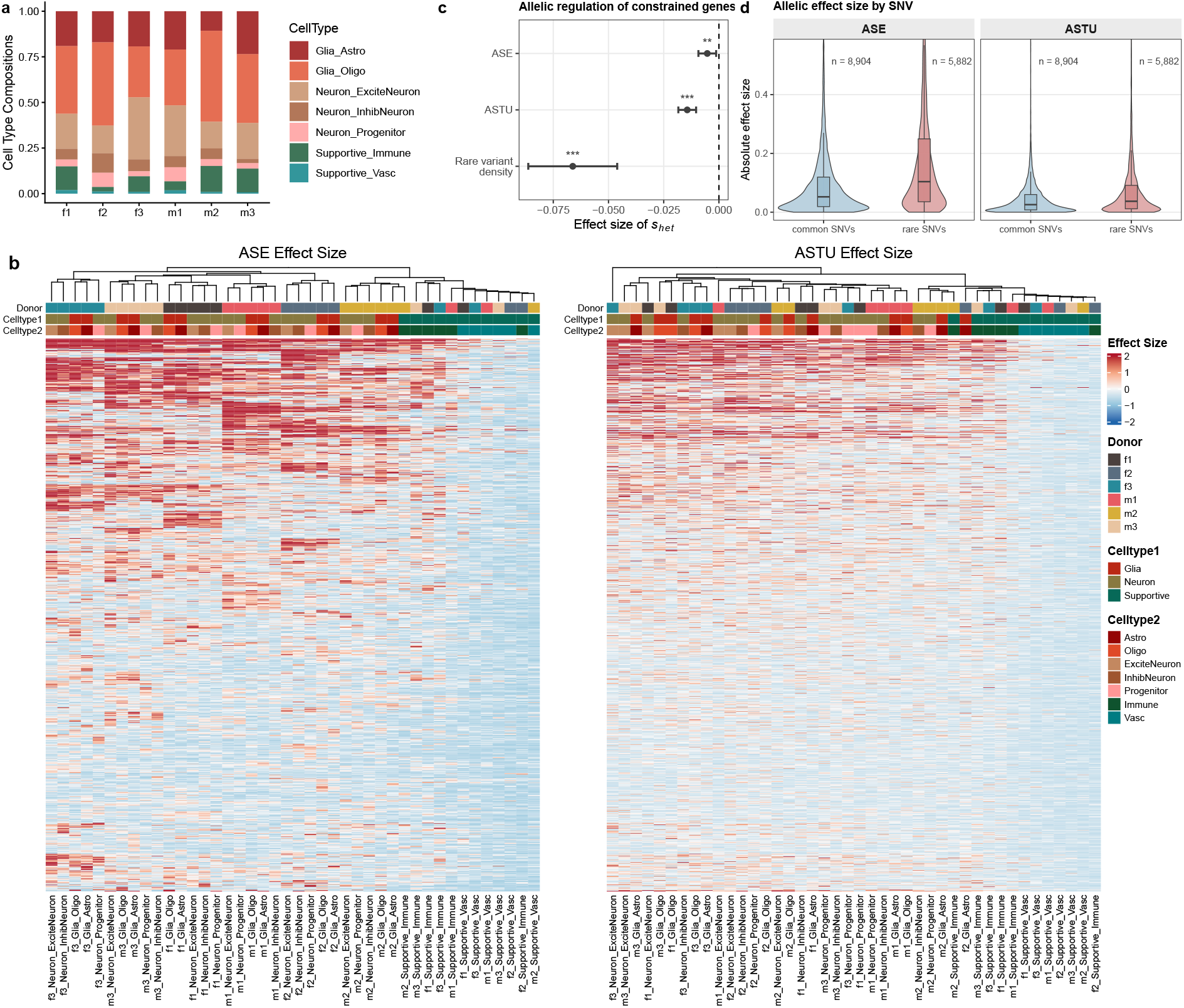
Allelic regulation under purifying selection in healthy human hippocampus long-read single-nucleus RNA-seq. LongAllele reveals that purifying selection constrains allelic imbalance at both gene-expression and isoform-usage layers and that rare variants exert disproportionately large allelic effects. (**a**) Stacked bar plot showing cell-type composition across 6 healthy donors and 7 cell types. (**b**) Hierarchically clustered heatmaps of ASE (left) and ASTU (right) effect sizes for the top 1,000 most variable genes. Columns (samples) cluster primarily by donor rather than cell type on both panels, with Supportive cells (Vascular, Immune) separating from Neuron/Glia subtypes in secondary branches. (**c**) Forest plot showing associations between the gene constraint score *s*_het_ (GeneBayes) and three allelic regulation metrics, ASE effect size (top), ASTU effect size (middle), and rare-variant density (bottom). Error bars represent 95% CIs of effect size estimates. ** *p <* 0.01, *** *p <* 0.001. Negative estimates across all three rows indicate that constrained genes show smaller allelic imbalance and fewer rare variants. (**d**) Absolute allelic effect size of ASE and ASTU between common and rare variants. Numbers above each violin indicate sample size. Violins show distribution density with inset boxes (median and quartiles). Rare variants show approximately 2-fold larger effects on both axes.

Beyond cell-type context, we next assessed how gene-level evolutionary constraint shapes allelic imbalance at both the gene-expression and isoform-usage axes. Using linear mixed-effect models across 10,893 phased autosomal genes (adjusted for cell type, gene expression and donor), we found that more constrained genes showed smaller ASE and ASTU effect sizes (*β*_ASE_ = −0.005, *p* = 9.9 × 10^−3^; *β*_ASTU_ = −0.014, *p* = 2.7 × 10^−12^, Fig. 5c). This pattern was consistent across alternative constraint metrics (LOEUF [52] and Przeworski *h*_*s*_ [53]; Table S6). This result is consistent with previous studies showing that constrained genes exhibit reduced cis-eQTL density and lower allelic imbalance [54]. The depletion of ASTU suggests that this selective constraint extends to the isoform-usage layer, in line with transcript-level mechanisms by which constrained genes can buffer variant effects through isoform choice [55].

Because rare variants often harbor the most pronounced allelic effects and serve as primary targets of purifying selection [15, 54], we further utilized LongAllele to characterize the rare-variant contributions of allelic regulation. We first asked whether rare variants in ASTU-significant genes are preferentially located in specific transcript structural elements. These rare variants were significantly enriched in intronic regions (OR = 1.83, *p*_adj_ = 3.7 × 10^−23^) and depleted in UTR regions (OR = 0.70, *p*_adj_ = 2.0 × 10^−6^) and synonymous variants (OR = 0.69, *p*_adj_ = 5.0 × 10^−5^) (Fig. S8). This pattern suggests that rare variants in ASTU-significant genes preferentially fall in annotations linked to transcript-structural regulation, such as intronic splicing regulatory elements, rather than in those expected to act primarily through protein and mRNA stability. To quantify rare-variant effects at the gene level, we focused on genes containing a single heterozygous SNV per donor, where the gene-level allelic effect directly serves as a proxy for that variant’s regulatory impact (Methods). In this setting, rare variants showed approximately 2-fold larger effects than common variants on both the ASE and ASTU axes (Wilcoxon rank-sum *p <* 10^−170^ for both; Fig. 5d), whereas the density of rare variants per gene declined significantly with increasing gene constraint (*p* = 1.2 × 10^−10^; Fig. 5c). Together, our results in the hippocampus demonstrate that purifying selection constrains allelic regulation at both gene and isoform levels, with constrained genes exhibiting smaller ASE and ASTU effect sizes and harboring sparser yet more impactful rare variants, extending classical purifying-selection signatures to haplotype-resolved isoform usage.

### 2.6 LongAllele resolves disease-associated regulatory effects in individual transcriptomes

To evaluate LongAllele in a disease-study setting, we applied it to long-read RNA-seq from dorsolateral prefrontal cortex in two Alzheimer’s disease (AD) case-control cohorts: the Aguzzoli Heberle (AH) cohort of six AD donors and six cognitively healthy controls profiled by Oxford Nanopore [56], and the ENCODE cohort of six AD donors and five controls profiled across PacBio and Nanopore [57]. Having demonstrated that ASE are more tissues- and cell-type-dependent than ASTU, we next asked whether a similar contrast appears across AD status. We compared genes with allelic imbalance restricted to either AD or control donors against genes with consistent significance patterns (Methods). ASE showed greater disease-context dependency than ASTU in both cohorts, with 62% versus 35% disease-context dependent calls in AH and a similar trend in ENCODE (Fig. 6a).

**Fig. 6.**
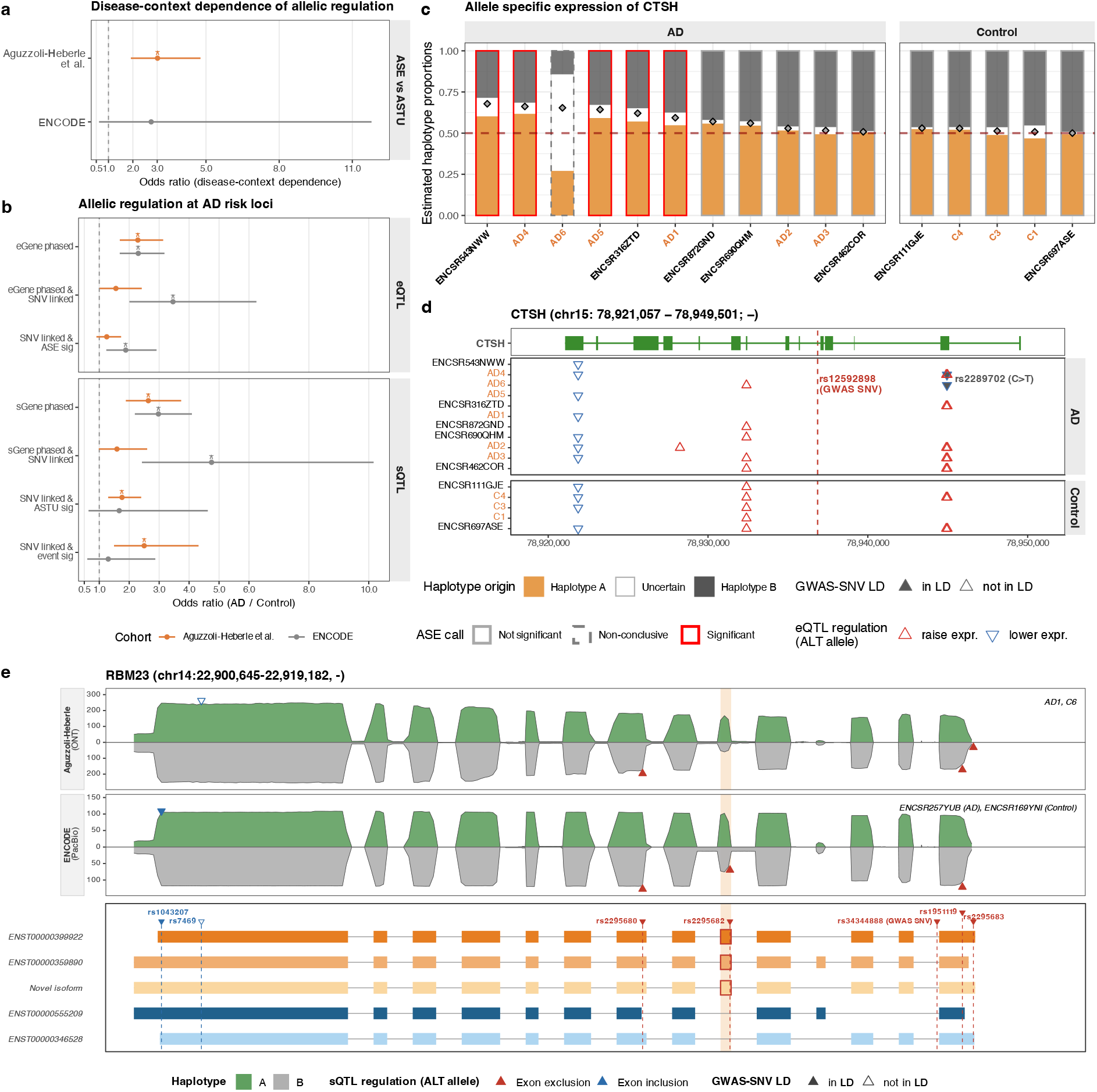
LongAllele supports variant interpretation in Alzheimer’s disease cohorts. (**a**) Disease-context dependency of ASE versus ASTU. Odds ratios compare how often ASE versus ASTU calls were significant in only AD or control donors within each cohort. (**b**) Recovery of AD-linked regulatory signals in AD versus control donors. For eQTL-linked eGenes and sQTL-linked sGenes, odds ratios compare AD donors with controls at three levels: genes phased by LongAllele, genes with a curated or LD-linked heterozygous SNV called, and genes with significant ASE, ASTU, or event-level HAEU/HAJU. (**c**) Haplotype-aware gene expression of *CTSH*. Bars show estimated haplotype proportions, with border colors indicating ASE call significance. (**d**) Per-donor heterozygous SNVs called by LongAllele and used to phase *CTSH*. The dashed line marks the intronic AD GWAS variant rs12592898, and rs2289702 is highlighted. Filled and open triangles indicate SNVs inside or outside the rs12592898 LD set, respectively; colors denote the eQTL direction of the alternative allele (red, higher expression; blue, lower expression). (**e**) Cross-platform resolution of an *RBM23* HAEU event. Haplotype-resolved coverage tracks show the event in AH Oxford Nanopore and ENCODE PacBio data, with the shaded region marking the cassette exon. Transcript models and triangles show the AD GWAS/sQTL variant rs34344888 and LD-linked SNVs; triangle colors denote the sQTL direction of the alternative allele (red, exon exclusion; blue, exon inclusion).

Given the limited cohort sizes, we focused on whether LongAllele can aid disease studies by resolving AD-associated regulatory effects in individual transcriptomes. We therefore curated two lists of AD-relevant variant-gene pairs by selecting AD GWAS [1] risk variants that are also reported as brain eQTLs or sQTLs in AD-relevant cohorts cataloged by NIAGADS FunGen-AD [58–62]. This curation yielded 3,269 eQTL-linked and 2,817 sQTL-linked variant-gene pairs, spanning 1,199 eGenes and 923 sGenes (Methods). We next asked how often these AD-linked variant-gene pairs could be resolved in individual transcriptomes (Fig. 6b). Across this analysis, AD donors generally showed greater recovery of disease-relevant signals than controls, from AD-linked eGenes and sGenes that could be analyzed by LongAllele, to genes in which a linked SNV was observed, to genes with significant allelic imbalance signals.

*CTSH* is a lysosomal protease gene in our AD-linked eGene set with established genetic and regulatory evidence for AD relevance. LongAllele detected significant ASE for *CTSH* in three AD donors from the AH cohort and two from the ENCODE cohort, whereas phased control donors remained allelically balanced (Fig. 6c). Population-level colocalization analyses indicate that AD risk and brain *CTSH* expression share a genetic basis at this locus [63]. LongAllele complemented this population-level evidence by resolving donor-level *CTSH* allelic expression through expressed heterozygous SNVs, including eQTL-linked variants called in phasable donors (Fig. 6d). Among the variants called by LongAllele, rs2289702 was an exonic missense SNV in LD with the intronic AD GWAS variant rs12592898 (r^2^ = 0.76; Fig. S9a) and previously implicated as a functional regulatory variant. LongAllele called rs2289702 in AD4, where *CTSH* ASE was significant, and in AD6, where the ASE call was inconclusive because many reads were non-phasable. In both donors, the ALT allele was associated with lower *CTSH* allelic expression. This direction was concordant with findings from Li et al., who identified rs2289702 through reporter, electrophoretic mobility-shift, and *ZFP69B* -knockdown assays, showing that the REF allele increases *CTSH* expression and confers AD risk through *ZFP69B* binding, whereas the protective ALT allele disrupts this effect [64].

We next examined *RBM23*, an RNA-binding protein gene among our AD-linked sGenes. LongAllele detected significant ASTU across several donors (Fig. S10), whereas gene-level ASE was largely balanced (Fig. S11). By resolving transcript-level imbalance into local splicing events, LongAllele identified a cassette-exon event with haplotype-biased inclusion across four donors (AD1, C6, ENCSR169YNI, and ENCSR257YUB), despite differences in donor genotype, disease status, and sequencing platforms. These donors were phased through different expressed heterozygous SNVs, most of which fell within the same LD block as the intronic AD GWAS variant rs34344888 (Fig. S9b), a reported brain sQTL for *RBM23* whose alternative allele is associated with cassette-exon exclusion [58, 59]. Across these donors, LongAllele resolved cassette-exon usage in the same direction as the brain sQTL (Fig. 6e), providing donor-level transcript-usage and event-level splicing evidence at an AD-associated sQTL. Together, these examples illustrate how LongAllele can support disease variant interpretation by adding donor-level expression and splicing evidence to population-level genetic associations across long-read platforms.

## 3 Discussion

In this study, we presented LongAllele, a statistical framework for joint haplotype inference and phasability-aware, multi-scale allelic testing from long-read bulk and single-cell RNA sequencing. The central methodological advance of LongAllele is to address limitations in both stages of allelic analysis. For haplotype inference, LongAllele jointly models heterozygous variant status, haplotype structure and read-haplotype assignment, rather than passing information through standalone variant callers and phasers. This formulation allows SNV evidence and haplotype structure to refine each other through within-read co-occurrence patterns, reducing error propagation into downstream allelic inference. In simulations and real datasets, this improved variant calling, haplotype phasing and haplotype-resolved quantification (Fig. 2, Fig. 3a; Table S1). For allelic testing, LongAllele explicitly accounts for non-phasable reads, whereas existing methods still discard them and leave their potential to bias allelic estimates unaccounted for, despite previous recognition of limited read phasability as a source of power loss [34]. By bounding allelic estimates under alternative assignments of non-phasable reads, LongAllele distinguishes significant, non-significant and inconclusive genes, reducing overconfident calls under phasability uncertainty (Fig. 2d, Fig. 3c). LongAllele further extends allelic testing across gene-level ASE, isoform-level ASTU and local haplotype-associated exon and junction usage (HAEU and HAJU) at both bulk and cell-type resolution, enabling more complete interpretation of haplotype-resolved variant effects from long-read RNA sequencing.

Across GTEx, PBMC, hippocampus and AD datasets, LongAllele consistently revealed greater context dependence of allelic regulation for gene expression than transcript usage (Fig. 3e, Fig. 4c, Fig. S7, Fig. 6a). One plausible explanation is that *cis*-regulatory effects on gene expression often act through context-dependent promoter-enhancer interactions shaped by transcription factor binding and chromatin accessibility [65, 66]. By contrast, allelic effects on isoform usage are more directly shaped by local sequence features, such as RNA-binding protein motifs [67], alternative transcription start sites [68] and polyadenylation signals [69], than by the broader *trans*-regulatory context. The same distinction is evident in our single-gene examples (Fig. 3f,g, Fig. 4b,d), where variant effects on isoform usage are largely consistent across tissues or cell types, whereas those on gene expression appear more context-dependent. In the hippocampus dataset, LongAllele further showed that purifying selection constrains allelic imbalance at both the gene and isoform levels (Fig. 5c). Specifically, more constrained genes showed smaller ASE and ASTU effect sizes with lower rare-variant density, whereas rare variants themselves tended to have larger allelic effects. Such a pattern is consistent with the action of purifying selection on dosage-sensitive genes, where intolerance to persistent allelic imbalance progressively depletes high-impact regulatory variants [15, 54, 55]. Collectively, these biological insights illustrate how haplotype-resolved analysis across gene, transcript and local-event levels can reveal regulatory patterns that would be difficult to detect from any single layer alone.

Despite these advances, LongAllele has several limitations that motivate directions for future work. First, the current framework assumes a fixed diploid genome shared by all reads and does not explicitly model cell-to-cell genetic heterogeneity arising from somatic mutations. Extending the model to capture such heterogeneity would enable applications in cancer and mosaic tissues. Second, LongAllele’s joint inference framework can become less stable in genes with limited haplotype phasing information, where spurious co-occurrence patterns from a large number of noisy SNVs can bury the true signal, leading to overconfident false calls (Fig. 3b,c). Although this risk is mitigated by carefully designed filters, including heterozygous-probability, homopolymer, RNA-editing and variant-cluster filters, as well as an optional XGBoost classifier for better handling sequencing artifacts in real data, a trade-off between sensitivity and robustness remains. Third, RNA-based genotyping is inherently restricted to transcribed regions, so causal distal non-coding variants may remain unobserved without paired genomic data. Although RNA-called SNVs could serve as sparse anchors for imputation [70, 71] or linkage-based prioritization of nearby regulatory variants, this strategy becomes less reliable over increasing genomic distance and is particularly limited for rare variants, which often carry the largest regulatory effects. Finally, LongAllele currently focuses on within-individual haplotype comparisons and gene-level phasing, and extending the framework to incorporate between-individual [2, 72] or longer-range [31] information will be an important direction for large-cohort applications.

In summary, LongAllele provides a unified and reliable framework for haplotype-resolved allele-specific analysis of transcriptional isoforms from long-read single-cell RNA sequencing. By jointly modeling variant calling and phasing with read phasability explicitly accounted for, it improves both the accuracy and reliability of allelic inference. These advances enable systematic characterization of *cis*-regulatory effects across gene, isoform and local-event levels in tissue, cell-type and disease contexts. We anticipate that this framework will be particularly useful for prioritizing candidate regulatory variants and for guiding experimental studies of variant effects across tissues, cell types and disease states.

## 4 Methods

### 4.1 Variant calling and SNV filtering

LongAllele takes as input a genome-aligned BAM file and performs variant calling through direct pileup, using read names from SCOTCH’s read-to-gene mapping as a per-gene whitelist. At each genomic position within a gene, only canonical bases (A, C, G, T) are counted toward depth and alternative allele support. Positions are retained if both total depth and alternative allele count exceed user-specified thresholds. When paired genotype data (e.g., from whole-genome sequencing) are available, the provided heterozygous SNV set is used directly and the filtering steps below are skipped. Otherwise, candidate SNVs are filtered through the following criteria sequentially by (1) heterozygous probability, (2) homopolymer context, (3) RNA editing sites, and (4) variant clustering.

(1) Heterozygous probability. For each candidate SNV, let *d* denote the total read depth and *a* the number of reads supporting the alternative allele. The posterior probability that the site is heterozygous is computed under a three-genotype model with genotypes *g* ∈ { het, hom-ref, hom-alt }, assuming a sequencing error rate *ϵ* (default *ϵ* = 0.01) and equal prior probability for each genotype. The genotype-specific likelihoods are

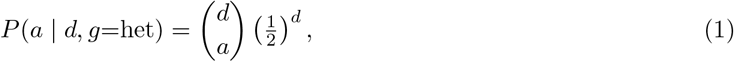

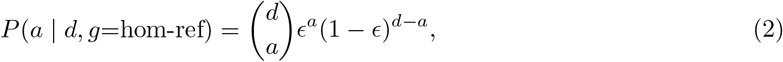

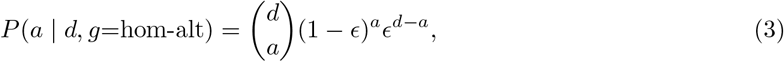

and the posterior heterozygous probability is

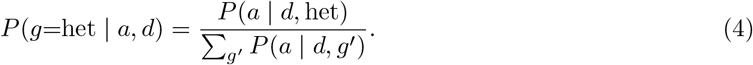

SNVs with *P* (*g*=het | *a, d*) *<* 0.99 are discarded (default threshold). The number retained per gene is further capped at ⌈6.6 × *L/*1000 ⌉ (*L*: total exonic length in bp), where 6.6 is the 97.5th percentile of heterozygous SNV density per kilobase estimated from the GIAB HG001 reference sample, retaining only the top-ranked candidates by heterozygous probability. For simulated data, where the candidate pool contains fewer spurious variants, an optional stepped fallback is available: if no SNVs pass the threshold, it is decreased by 0.05 iteratively down to a floor of 0.5, and the first passing set is retained (capped at the same density limit). This fallback is disabled by default for real data, where high sequencing depth causes homozygous-alternative and error sites to accumulate sufficient alternative allele support to pass depth thresholds, and the strict filter appropriately excludes such sites. (2) Homopolymer filter. SNVs located near homopolymer regions are removed. A window of ±20 bp around each candidate is examined, and SNVs adjacent to homopolymer runs (≥ 5 consecutive identical bases or repeat k-mer units) are flagged due to elevated indel error rates in long-read sequencing. Flagged SNVs are discarded unless they show strong alternative allele support (alt count ≥ 20 by default), preserving true variants in repetitive contexts. (3) RNA editing filter. A*>*G and reverse-complement T*>*C substitutions matching known adenosine-to-inosine RNA editing sites in REDIportal v3 [36] are then removed to avoid misidentification of RNA editing events as genomic variants. (4) Variant cluster filter. Clustered SNVs are filtered out by removing regions with high variant density (≥ 3 SNVs within a 20bp window by default), which often reflect alignment artifacts or complex variation. As above, SNVs with strong alternative allele support (alt count ≥ 20 by default) are retained to avoid discarding true variants in polymorphic regions.

To improve EM convergence in the presence of context-dependent artifacts, LongAllele trained an XGBoost classifier [37] on Genome in a Bottle (GIAB) HG001 (NA12878) Oxford Nanopore cDNA RNA-seq data [20] to score each candidate as an informative initialization for the SNV inclusion parameter 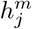. The classifier includes 17 features spanning four categories: read-level statistics (depth, alternative allele count, posterior heterozygous probability), base and mapping quality (mean mapping quality, mean base quality for alternative and reference alleles), strand and positional bias (strand odds ratio, alternative allele position mean and standard deviation on reads, number of distinct alternative alleles), and sequence context (GC content in an 11 bp window, homopolymer run length, whether the substitution creates a new homopolymer or occurs in an AT-rich flanking context, and transition versus transversion status). Candidates scoring below a user-specified threshold (default 0.05) may additionally be removed as obvious artifacts prior to EM.

Single-nucleus RNA-seq libraries can contain substantial unspliced pre-mRNA, often referred to as nascent-RNA leak. In long-read snRNA-seq, intron-dominated reads can have ambiguous isoform origin and introduce artifacts into haplotype phasing. To mitigate this issue, LongAllele provides an optional high-artifact mode (`--high artifact mode`) with two disabled-by-default filters. At the SNV level, LongAllele uses both exonic and intronic SNVs as phasing markers by default, but in high-artifact mode excludes SNVs in read-dense intronic regions for high-leak genes (intronic territory broadly covered by reads *>* 0.25) to prevent nascent intronic allelic biases from distorting read-haplotype assignment for mature transcripts. At the read level, reads whose alignments lie predominantly within introns (default fraction *>* 0.60) are also excluded. These filters are intended for snRNA-seq datasets where nascent-RNA leak is a concern.

### 4.2 EM algorithm for joint variant calling and haplotype phasing

LongAllele performs joint inference of variant calls and haplotype phasing via the expectation-maximization algorithm [73]. For a given gene, let *N* denote the number of reads and *M* the number of surviving candidate heterozygous SNVs after filtering. The observation at the *j*^*th*^ SNV for read *i* is encoded as *r*_*ij*_ ∈ { ref, alt, other }, with position-specific sequencing error rate *π*_*ij*_ ∈ [0, 1] derived from base quality (Phred) scores. Note that, reads not covering a given SNV position are treated as missing, and thus do not contribute to the likelihood function. The model introduces two sets of latent variables: *I*_*i*_ ∼ Bernoulli(*α*_*A*_), representing whether read *i* originates from haplotype A, and 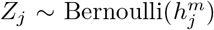, representing whether SNV *j* is truly heterozygous (*Z*_*j*_ = 1) rather than sequencing noise (*Z*_*j*_ = 0). The parameters to be estimated are the gene haplotype-A proportion *α*_*A*_ (with *α*_*B*_ = 1 − *α*_*A*_ and *α*_min_ = min(*α*_*A*_, *α*_*B*_)), the SNV heterozygosity probability *h*^*m*^, and the haplotype-A mapping probability 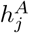 for each SNV, with 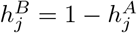.

The complete likelihood function factorizes as:

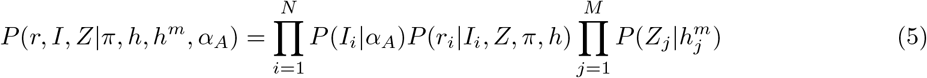

The per-read, per-SNV likelihood is:

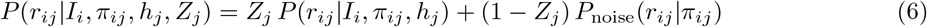

where

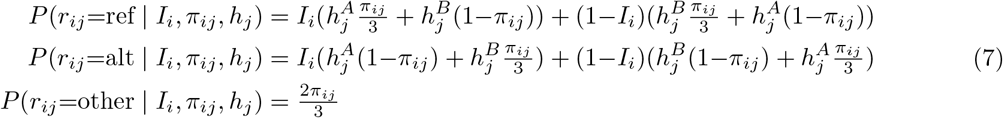

and

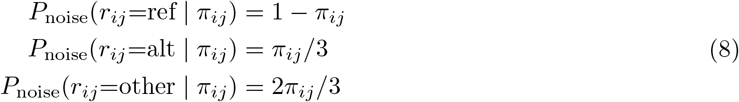

In the E-step, the posterior haplotype probability for each read is:

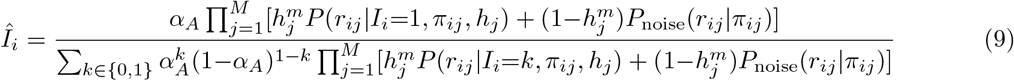

and the posterior heterozygosity probability for each SNV is:

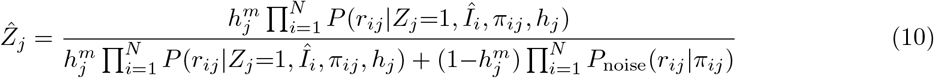

In the M-step, *α*_*A*_ and *h*^*m*^ have closed-form updates:

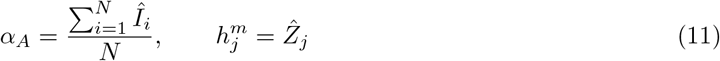

The haplotype-A mapping probability *h* is updated by numerically maximizing the expected complete log-likelihood. For each SNV *j, h*_*j*_ maximizes

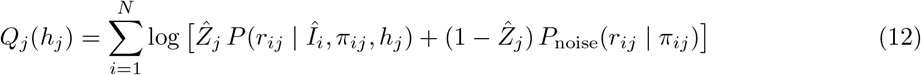

which has no closed-form solution and is solved numerically. The algorithm iterates until either the maximum parameter change or the change in the expected complete log-likelihood falls below a convergence threshold *ϵ*.

To avoid convergence to local optima of the EM algorithm, LongAllele initializes the haplotype-A mapping probability *h* using spectral clustering on a SNV co-occurrence matrix, where each entry counts the number of reads carrying the alternative allele at both SNVs *j* and *j*^*′*^. The two major clusters from spectral clustering are assigned as haplotype A and haplotype B, with corresponding 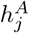 initialized at 0.9 and 0.1, respectively. SNVs that are less connected to other SNVs within their cluster are initialized as 0.5 with a small perturbation, allowing the EM to resolve their haplotype mapping from data. The initial *α*_*A*_ is set to 0.5. Each 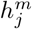 is initialized as the posterior heterozygous probability under the same three-genotype model used in SNV filtering, but with per-read base quality scores set as *π*_*ij*_ instead of being fixed. Alternatively, 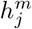 can be initialized from external sources such as a trained variant classifier or user-supplied confidence scores, which may improve convergence when sequencing artifacts inflate the candidate pool. The EM re-estimates 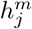 during inference regardless of initialization source.

### 4.3 Detection of allelic imbalance

LongAllele tests for ASE using a likelihood ratio test (LRT) with hypotheses:

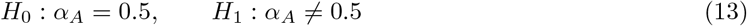

The observed log-likelihood marginalizes over both latent variables *I* and *Z*. Defining the per-read, per-SNV marginal as

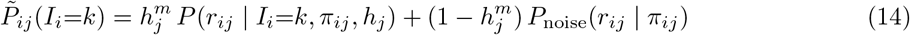

the observed log-likelihood is:

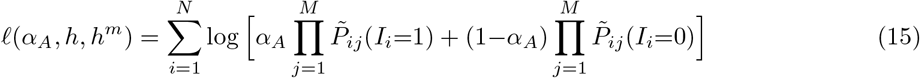

At the bulk level, the alternative log-likelihood uses the EM estimates, while the null log-likelihood re-estimates *h* and *h*^*m*^ with *α*_*A*_ fixed at 0.5 via a constrained EM:

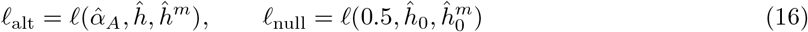

The test statistic follows a 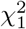 distribution under *H*_0_:

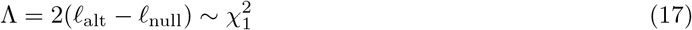

At each cell type, LongAllele re-runs the EM algorithm on corresponding read subset with *h* and *h*^*m*^ being fixed at the bulk-level estimates (*ĥ, ĥ*^*m*^) and updating only *α*_*A*_ and the read-haplotype posteriors *Î*. This ensures consistent variant calls across cell types while allowing cell-type-specific haplotype proportions. The cell-type alternative and null log-likelihoods are:

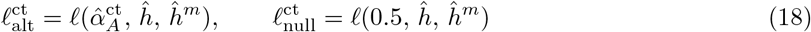

LongAllele tests for ASTU using a chi-squared test of independence on a *K ×* 2 contingency table, where *K* is the number of isoforms and the two columns correspond to haplotype A and haplotype B. Each cell contains the sum of posterior haplotype probabilities (*Î*_*i*_ or 1 − *Î*_*i*_) across reads assigned to that isoform by SCOTCH, yielding fractional weighted counts. The hypotheses are:

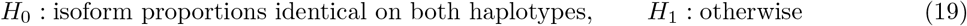

Isoforms with expression fraction below a minimum threshold (default 10%) on both haplotypes are collapsed into an other category to avoid inflating degrees of freedom with rare isoforms. The test statistic follows a 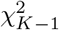 distribution under *H*_0_, where *K* is the number of categories after collapsing. At the cell-type level, the ASTU test subsets the bulk-level haplotype-weighted count matrices by cell type, rather than re-estimating haplotype posteriors per cell type.

LongAllele also reports an effect size per gene to quantify the magnitude of allelic imbalance. For ASE, the effect size is the log2 ratio of read counts between the major and minor haplotypes, derived from 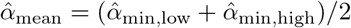 and stabilized by a shrinkage pseudocount *k* set adaptively to 5% of the median gene-level read count. For ASTU, the effect size is the log2 ratio of the dominant-isoform usage fractions between haplotypes, with shrinkage scaled sub-linearly by 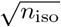 to maintain calibration across multi-isoform genes.

To facilitate variant interpretation, LongAllele identifies haplotype-associated exon usage (HAEU) and haplotype-associated junction usage (HAJU) through chi-squared tests among all phased reads of the gene. Here, LongAllele uses read-isoform mappings and transcript structure annotations from SCOTCH for testing rather than directly from BAM alignments, to avoid alignment and sequencing artifacts. Candidate SNVs are then linked to each event if they lie within a preset exonic distance (± 50 bp by default), and potential regulatory direction can be inferred from the SNV-haplotype mapping probabilities, read-isoform assignments, and read-haplotype origin probabilities. LongAllele also tests SNV-event pair associations using chi-squared tests among all reads covering the variant site within the gene, regardless of haplotype origin. Optionally, haplotype-event testing can be restricted to events at isoform-switching boundaries, defined by the structural differences between the most decreased and most increased isoforms between haplotypes.

### 4.4 Phasability-aware significance calling

Because not all reads overlap a heterozygous SNV, allelic tests based solely on phasable reads may yield overconfident or underpowered results. LongAllele addresses this by computing bounds that reflect the uncertainty introduced by non-phasable reads, and reports each gene as significant, non-significant, or inconclusive.

For ASE, bounds on the minor haplotype proportion estimations 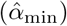 are computed by assigning all non-phasable reads to one haplotype or the other, yielding 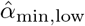 and 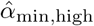 after relabeling to the minor haplotype. A gene is called significant ASE if *p*_adj_ ≤ 0.05 and 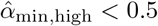, confirming that allelic imbalance persists even under the most conservative assumption about non-phasable reads. A gene is non-significant if *p*_adj_ *>* 0.05, and inconclusive otherwise.

For ASTU, three versions of the isoform-by-haplotype contingency table are constructed and each tested using a chi-squared test of independence: one using only phasable reads (*p*_adj_), one where non-phasable reads are assigned their mean posterior haplotype probability from the phasable subset (tending toward balanced allocation, yielding the most conservative *p*_adj,high_), and one that reallocates non-phasable reads to maximize imbalance between haplotypes (yielding the most sensitive *p*_adj,low_). A gene is called significant ASTU if *p*_adj,high_ ≤ 0.05, non-significant if *p*_adj,low_ *>* 0.05, and inconclusive otherwise.

### 4.5 Allelic cell-type variability

To quantify the extent to which allelic regulation varies across cell types, we defined allelic cell-type variability (ACTV) to measure how much a gene’s allelic regulation varies across cell types within each donor. For a gene observed in cell types CT_1_, CT_2_, …, CT_*k*_, we computed a haplotype gene expression difference *δ*_ASE_ and a haplotype isoform usage difference *δ*_ASTU_ in each cell type. For ASE:

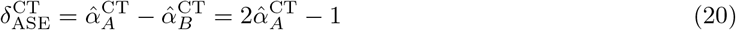

where *δ*_ASE_ measures the degree and direction of haplotype-A dominance in gene expression. For ASTU:

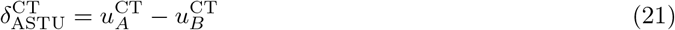

where *u*_*A*_ and *u*_*B*_ are the usage fractions of a representative isoform (selected as the isoform with the largest |*δ*_ASTU_| in bulk) on haplotypes A and B, respectively, and *δ*_ASTU_ measures the degree and direction of haplotype-specific isoform preference. ACTV is then defined as the range of *δ*_ASE_ and *δ*_ASTU_, respectively, across cell types:

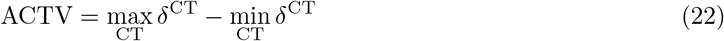

where *δ* ∈ { *δ*_ASE_, *δ*_ASTU_ }. Larger values of ACTV indicate stronger cell-type dependence, with values exceeding 1.0 necessarily involving direction reversal across cell types, while ACTV = 0 indicates identical allelic effects across all cell types (constitutive regulation). ACTV computation was restricted to genes with ≥ 50 phasable reads per cell type in at least two cell types and called significant by the phasability-CI-robust criterion (Methods 4.4) in at least one cell type for the respective metric. Genes not passing the significance call were assigned ACTV = 0.

### 4.6 Simulation setup and benchmarking

We estimated per-gene heterozygous SNV densities from the Genome in a Bottle (GIAB) HG001 reference sample [20] on chromosome 6, yielding 3,627 SNVs across 464 genes (1 to 38 heterozygous exonic SNVs per gene, with a median of 6), and randomly allocated each SNV to one of two haplotypes in equal proportion. Per-gene read counts were sampled from *n*_*g*_ ∼ LogNormal(5.62, 1), with parameters estimated from a long-read scRNA-seq PBMC dataset [35], and reads were allocated to haplotype A via *n*_*A*_ ∼ Binomial(*n*_*g*_, *α*_min_) with *α*_min_ ∈ {0.1, 0.3, 0.5}, with the remainder assigned to haplotype B. Isoform proportions for each haplotype were sampled from ***π***_*h*_ ∼ Dirichlet(***α***_*h*_), *h* ∈ { *A, B* }. For 160 ASTU genes, the two haplotypes received concentration parameters from different distributions (Gamma(2, 2) and Gamma(0.5, 0.5)) to induce distinct isoform compositions. For the remaining 304 non-ASTU genes, ***α***_*A*_ = ***α***_*B*_ = ***α*** with ***α*** ∼ Gamma(2, 2). Simulated FASTA reads were generated using the LRGASP simulation pipeline [74] and aligned to the GRCh38 reference genome with minimap2 [75]. Isoform structures were defined using GENCODE v40 annotation. We simulated two phasability conditions. In the fully phasable setting (464 genes), every isoform structure contains at least one exonic heterozygous SNV, so that the vast majority of reads are phasable, with a small fraction remaining non-phasable due to sequencing and alignment artifacts. In the partially phasable setting (376 multi-isoform genes), at least one isoform per gene does not cover any exonic heterozygous SNV, producing non-phasable reads. LongAllele was run without paired genotype data, using the statistical heterozygous probability as 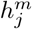 initialization. LongAllele-Genotyped received ground-truth SNV sites as input.

For haplotype proportion estimation and ASE testing, isoLASER’s 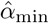 was computed from the largest bi-haplotype phasing group per gene, and HAPCUT2’s from phased SNV-level read counts, with ASE significance assessed via binomial tests in both cases. longcallR provides native ASE quantification. Since isoLASER, longcallR, and HAPCUT2 do not provide a direct test for haplotype-specific isoform composition changes, we evaluated ASTU detection using each method’s closest available output. Specifically, we used isoLASER’s mutual information filtering on haplotype-associated splicing events, longcallR’s allele-specific junction (ASJ) gene-level test, and for HAPCUT2, a chi-squared test of independence on isoform-by-haplotype contingency tables constructed from its phased read assignments and SCOTCH isoform annotations. All methods were run with default parameters. We additionally evaluated longcallR with its dense-SNP cluster filter disabled (longcallR, no dn filter) to isolate the effect of this filter on variant calling performance.

### 4.7 Data processing and statistical analysis

All real datasets were sequenced on the Oxford Nanopore Technology (ONT) platform and processed through SCOTCH [35] and LongAllele sequentially. GTEx v9 long-read bulk RNA-seq data (seven donors, each profiled in two tissues) and paired whole-genome sequencing genotypes were obtained from the GTEx Portal [2, 25]. LORALS ASE results were downloaded from the published output [25]. LORALS does not provide ASTU analysis. Linkage disequilibrium was computed using LDlinkR [76] with 1000 Genomes Phase 3 [77] reference data (GRCh38, all populations). Long-read single-cell RNA-seq data of PBMCs (two donors) and cell type annotations were obtained from [35]. Long-read single-nucleus RNA-seq data of human hippocampus (six healthy donors, three female and three male) and cell type annotations were obtained from [51]. Long-read bulk RNA-seq of human dorsolateral prefrontal cortex was obtained from two AD case-control cohorts: the Aguzzoli Heberle cohort (six AD and six control donors, Oxford Nanopore) [56] and the ENCODE cohort (six AD and five control donors, PacBio and Oxford Nanopore) [57], after excluding one low-coverage ENCODE control donor (ENCSR094NFM).

For the heatmap visualization in Fig. 5b, we restricted to the top 1,000 most variable genes (by cross-sample × cell-type variance) with bulk phasable depth ≥ 100 across all donors. For the constraint-dependence analysis in hippocampus, linear mixed-effects models were fit to z-scored ASE and ASTU effect sizes as a function of the gene constraint score *s*_het_ (log-transformed, z-scored), adjusting for cell-type group (Neuron or Supportive), read depth (linear and quadratic terms), and a donor random intercept. Analyses were restricted to (gene, cell-type, donor) triples, excluding bulk aggregates. Models were fit by REML using `lme4` [78], and fixed-effect significance was assessed by likelihood-ratio tests following ML refitting with `lmerTest` [79]. Absolute ASE and ASTU effect sizes were compared between rare and common variants using two-sided Wilcoxon rank-sum tests, restricting analyses to genes with exactly one heterozygous SNV per donor. For enrichment analysis of rare-variant functional classes, rare variants (maximum gnomAD v4.1 allele frequency *<* 0.01 or absent) in ASTU-significant genes were compared with all rare variants across the six donors by VEP-annotated functional class [80]. Foreground variants were further restricted to those for which the only donor carrying the variant showed significant ASTU, while all other donors expressing the gene showed non-significant ASTU. Class-specific odds ratios and *p*-values were computed using binary logistic mixed-effects regression with a donor random intercept (`lme4` [78]) and read-depth covariates, with Benjamini-Hochberg adjustment across different functional classes. Rare-variant counts per gene were fit against *s*_het_ (log-transformed, z-scored) using negative binomial regression, with log-transformed exonic length as an offset and log-transformed mean bulk read depth (with one added) as a covariate to adjust for gene expression. Significance of *s*_het_ was assessed by a Wald test [81]. Two AD-relevant variant-gene pair lists were curated by intersecting AD risk variants from the GWAS Catalog (“Alzheimer” trait, GRCh38) [1] with significant brain eQTL and sQTL associations (FDR *<* 0.05) from NIAGADS FunGen-AD [58], including ROSMAP [59], MSBB [60], Knight-ADRC [61] and MiGA [62]. For each curated variant-gene pair, LD-linked SNVs (*r*^2^ ≥ 0.6) were identified using 1000 Genomes phase 3 European-ancestry samples through LDlink [76]. Genes were evaluated in each donor at three nested levels: phased by LongAllele, phased with a curated or LD-linked heterozygous SNV called, and significant by the corresponding LongAllele allelic test. AD and control donors were compared using one-sided McNemar tests on discordant gene-level calls, with odds ratios reported as AD versus control. Disease-context dependency was assessed among genes with conclusive calls in at least three AD and three control donors. Genes significant in at least two donors in only one group were compared with genes showing consistent significance patterns, and ASE versus ASTU rates were tested by Fisher’s exact test. All statistical analyses in this section were performed in R v4.3.1 [82].

## Data availability

GTEx v9 long-read RNA-seq data and paired whole-genome sequencing genotypes were obtained from the GTEx Portal (https://gtexportal.org). Long-read single-cell RNA-seq data of PBMCs are available from [35]. Long-read single-nucleus RNA-seq data of human hippocampus are available from [51].

## Code availability

The LongAllele software is available at https://github.com/WGLab/LongAllele.

## Acknowledgements

We thank the IDDRC Biostatistics and Data Science core (NIH grant HD105354) for technical support on machine learning and high-performance computing. This study is supported by NIH grant HG013359 and the CHOP Research Institute.

## Author contributions

Z.X. conceived the study, developed the statistical framework, implemented the LongAllele software, performed all analyses, prepared the figures, and wrote the manuscript. K.W. supervised the project, acquired funding, and revised the manuscript. Both authors read and approved the final manuscript.

